# GANGE: Achieving Sequencing Without Sequencing With Diffusion Guided Generative Genomic Transformer

**DOI:** 10.64898/2026.04.15.718133

**Authors:** Sagar Gupta, Anchit Kumar, Umesh Bhati, Ravi Shankar

**Affiliations:** Studio of Computational Biology & Bioinformatics, The Himalayan Centre for High-throughput Computational Biology, (HiCHiCoB, A BIC supported by DBT, India), Biotechnology Division, CSIR-Institute of Himalayan Bioresource Technology (CSIR-IHBT), Palampur (HP), 176061, India; Academy of Scientific and Innovative Research (AcSIR), Ghaziabad-201002, India

**Keywords:** Genome assembly, Deep Learning, DDPM, Transformers, Genomics

## Abstract

The genome of a species is its book of life, but opening that book remains a costly affair due to the limitations the existing sequencing technologies pose. Short reads sequencers struggle to capture long and complex genomes, though have high fidelity rate. To counter that long reads from III^rd^ generation sequencers are used, which are full of indel errors. Thus, reads from both approaches are collectively used with very high coverage, making the sequencing projects unreasonably high of cost and unapproachable to majority. Here we present a first of its kind generative deep-learning system, GANGE, which not just recovers the correct sequence with high accuracy from indel prone ONT reads at manifold lesser coverage but also extends it by 4kb, achieving sequencing without sequencing, horizontally as well as vertically while maintaining >92% accuracy consistently. This all makes it possible to drastically pull down sequencing project cost. GANGE was tested across *A. thaliana*, *O. sativa* genomes and Human chromosome 1 where it delivered outstanding assembly performance. Besides this, it was also used to accurately generate 2kb upstream promoters of all the genes from 12 different species, demonstrating that one can now also take up regulomics research just using RNA data alone when genome sequence is not available. With this all, GANGE brings a democratic turning point in the area of genomics and sequencing research.

## Introduction

The field of genomics has been revolutionized by the advent of high-throughput sequencing (HTS) technologies which enabled researchers to decode the molecular life of several organisms with high speed and accuracy. Genome assembly, the process of reconstructing the original genome sequence from fragmented sequencing reads, is a cornerstone of any genomics research **[1]**. This can be achieved either through reference-guided assembly, or through *de novo* assembly where the genome is assembled directly from scratch without any prior reference. To sequence a new genome, which is mostly the case one has to take the second way. However, *de novo* assembly is highly demanding and expensive pursuit due to matters like lack of guide, demand of high fidelity reads, very high coverage, needs to counter complex repeats, pushing the sequencing project’s cost to unreasonable heights. More so, with length and complexity of genome.

For a 1 GB genome, Illumina sequencing costs at least $450-$800 for 30x coverage. Though this requires high-volume batching and results in highly fragmented assemblies that often fail to resolve structural complexity. In contrast, long-read platforms like Pacific Biosciences (PacBio) and Oxford Nanopore Technology (ONT) provide the structural integrity required for reference-grade genomes but at a higher premium. A 1 GB genome at 30x coverage on a PacBio system costs approximately $950-$1250, while ONT provides a more flexible price point of $800-$1250 with the unique advantage of spanning ultra-long repetitive regions that are otherwise unmappable. As genome size increases, the difficulties and associated costs scale non-linearly.

Second-generation sequencing (SGS) platforms, such as Illumina, produce highly accurate reads with error rates below 0.01% **[2]** but are limited to lengths of approximately 150-300 base pairs (bp) **[3]**. These short reads are suitable for high-throughput applications but often fail to resolve complex repetitive regions, structural variants, or long insertions and duplications. For genome sequencing they essentially require several other supporting approaches like mate-pair libraries, mapping guide, and very high coverage. The long-reads from the third generation sequencers have become their essential partners **[4]**. The third-generation sequencing (TGS) technologies, including PacBio **[5]** and ONT **[6]**, overcome many of these limitations by producing long reads with claimed average lengths of 10 kilobases (Kbp) or more, which are several orders of magnitude longer than SGS reads **[7–10]**. These long read lengths were supposed to enable the disambiguation against the repetitive regions and surpass them to reach the unique regions to obtain correct and longer assemblies **[11–12]**. Consequently, long-read sequencing has become a default component in *de novo* genome assembly **[13–15]**.

However, the benefit of increased read length is outweighed by the kind and level of sequencing errors. Both PacBio and ONT platforms exhibit high raw error rates, typically ranging between 15-25% with most errors resulting from indels, changing the sequence code enormously. The error distribution differs between the platforms **[15, 16–18]**. PacBio employs single-molecule real-time (SMRT) sequencing **[5]**, generating two major types of reads: **a)** Continuous Long Reads (CLR), which can exceed 25 kb in length but suffer from relatively high error rates (>15%). **b)** Circular Consensus Sequencing (CCS) or HiFi reads, introduced in 2021, which sequences the same molecule several times to achieve high accuracy (up to 99.99%) with read lengths between 10-20 kb, a coverage increasing strategy to counter the errors. ONT **[6]**, on the other hand, uses a nanopore-based system in which DNA molecules pass through protein nanopores, generating electrical signals interpreted into base sequences. ONT can produce ultra-long reads, sometimes surpassing 100 kb, which is highly advantageous for resolving structural variations and complex genomic regions. Among all the sequencers, it is the most affordable and easily accessibly one. In yet, long reads from both the platforms have very high error rates, making it necessary to neutralize them using high coverage and short reads from Illumina sequences, supplemented with various error correction algorithms in order to assemble them.

In this context, both traditional and machine learning (ML)-based error correction methods play an important role in ensuring the proper assembly of genomic data. A typical hybrid assembly pipeline proceeds in two main stages: **a)** Initial Assembly: Long-read assemblers, such as Canu **[19]**, Flye **[20]**, or HGAP **[21]**, generate a draft assembly using overlap-consensus or string graph-based methods. **b)** Polishing: Short reads (Illumina) are mapped to the draft assembly using tools such as BWA-MEM **[22]** or Bowtie2 **[23]**, followed by correction/polishing with tools such as Pilon **[24]** and POLCA **[25]**. For Nanopore data, ONT’s inbuilt software Nanopolish operates directly on the raw electrical signals, enabling it to refinement of basecalls and correction of systematic biases inherent to ONT technology.

One can also introduce error correction once these platforms release reads/super reads, using Illumina short reads. Error correction algorithms serve as the foundation for refining raw sequencing reads before assembly or polishing steps. They are the most critical determinant towards the assembly quality. These algorithms are designed to identify and fix sequencing errors mismatches, insertions, deletions, and structural misreads while preserving authentic genomic signals. Broadly, long-read error correction methods are broadly classified into self-correction and hybrid correction approaches. Self-correction relies exclusively on overlaps among long reads to generate consensus sequences, requiring very high sequencing depth and substantial computational resources (CONSENT **[26]** and VeChat **[27]**). On the other hand, hybrid correction integrates high depth of accurate short reads to correct long reads which include alignment-based (PacBioToCA **[28]**, Proovread **[29]**, Nanocorr **[30]**, and CoLoRMap **[31]**), assembly based (ECTools **[32]**, and HALC **[33]**), de Bruijn graph (DBG)-based (LoRDEC **[34]**, Jabba **[35]**, and FMLRC **[36]**), and hidden markov model (HMM)-based strategies (Hercules **[37]**), each offering different trade-offs between accuracy, computational efficiency, and scalability for correcting noisy long-read sequencing data. A common phenomenon observed is that the short reads give up correction if several indels/mismatches are encountered, as most of them utilize BWT based almost exact alignment with limited mismatch.

With the advent of machine and deep learning (ML/DL) approaches, this landscape has substantially changed by enabling models to learn directly from empirical data distributions and adaptively recognize complex, non-linear relationships between sequencing signals and true genomic sequences **[38]**. Traditional algorithms capture only short-range dependencies while deep-learners can represent high-dimensional and long-range dependencies without relying on restrictive probabilistic simplifications, making them particularly suitable for the noisy, high-dimensional nature of sequencing data **[39]**. Among the early ML-based error correction methods, Hercules **[37]** integrates the conceptual framework of HMMs with a learning-based parameter refinement process. Following Hercules, the field witnessed a paradigm shift toward artificial neural networks (ANNs), recurrent neural networks (RNNs), and, more recently, transformer-based architectures **[40]**, all of which excel in sequence modeling tasks. Transformers have since been adapted to genomics with impressive success. By treating nucleotide sequences in a manner similar to text data, transformer models **[41–44]** can learn the “grammar” of DNA. When trained on large-scale datasets, these models can accurately differentiate between systematic sequencing noise and true biological variations, making them ideal candidates for artificial intelligence (AI)-driven error correction in sequencing data.

DL has now been spread to nearly every stage of the sequencing and assembly pipeline **(Table S1)**. In the context of ONT, models such as DeepNano **[45]**, Chiron **[46]**, DeepNano-blitz **[47]** and Fast-Bonito **[48]** have employed convolutional and RNNs to translate electrical signals into base sequences, achieving a substantial increase in base-level accuracy. DL models like HECIL **[49]**, RNNHC **[50],** DeepCorr **[51],** and nmTHC **[52]** tried to solve error correction in long reads with hybrid approach. DeepPolisher **[53]** utilizes encoder-only transformers (EoT) to achieve highly accurate consensus sequences from PacBio data. For ONT reads, Herro **[54]** combines convolutional neural network (CNN) and EoT. **Table S1** highlights the various categories of the algorithms employed to every stage of the sequencing and assembly pipeline.

In addition to error correction, the significant advantages of TGS in producing reads that span kilobases, these sequences are frequently interrupted by biochemical damage or stochastic sequencing noise generating the kind of error which is usually difficult to deal: indels. They completely even change the sense of the sequence and disrupt contigs formation. When reads are fragmented, assemblers struggle to resolve “dead ends” in the assembly graph, leading to highly fragmented contigs and collapsed repetitive elements. In these conditions, extension of reads and resequencing is the solution. In past, read extension was attempted only through overlap-based extension, where reads were merged solely based on shared *k*-mer sequences. Tools like SSPACE or PBJelly used paired-end information or long-read scaffolds to bridge gaps **[55–56]**. However, these methods are often “reactive”, as they can only fill a gap if a physical read already exists to span it. All of them essentially require additional steps of long-reads sequences, mate pair library sequencing, and physical/digital mapping, raising the cost and complexities many notches more. They lack any “generative” capacity to handle horizontal coverage gaps or regions where the sequencing chemistry fails to produce a viable signal.

Thus, despite all this progress, sequencing any eukaryotic or big complex genome remains a highly costly affair. It essentially requires lots of sequencing and coverage to attain anything reliable. However, the kind of revolution generative AI is bringing, remains still largely untamed to demonstrate its full potential in resolving genome sequencing and issues. Recently, we developed generative deep learning systems which can perform generative tasks like condition-specific DNA methylation states of promoters **[57]** or *ab initio* transcription factor binding site (TFBS) discovery using just DNA sequence data **[58]**. Both these software tools has the capability to profess sequences which gives us hope to attempt generating sequences based on existing sequence data. It is possible to learn the noise in reads and generate the right bases there. And it is also possible to generate a whole sequence of bases while learning through the context foundation provided by the sequenced reads to extend it further into longer reads. Something like sequencing without sequencing!

Building upon the above possibility, we here introduce an innovative generative deep-learning framework termed **G**enerative **A**dditive **N**ucleotides based **G**enome **E**volver **(GANGE),** designed to perform reads error correction and their extension, while being capable to work at as low as 4x coverage for genome assembly. GANGE introduces a DL architecture that seamlessly integrates the Denoising Diffusion Probabilistic Model (DDPM) with a transformer-based encoder-decoder framework, effectively combining the stochastic generative power of diffusion models with the contextual learning and long-range dependency detection capabilities of transformers. This architecture enables GANGE to generate highly accurate nucleotide sequences (>92%) and reconstruct genomic sequences with unprecedented precision and generalization capacity. It achieves this all at much lesser coverage (as low as 4x), which the existing protocol are able to achieve only at 6 times or higher coverage. The system was trained on an extensive and diverse datasets comprising over 20 million genomic sequences, covering 20 sequence data, sourced from a broad spectrum of species. This large-scale, heterogeneous dataset allowed GANGE to learn universal genomic representations, facilitating reliable cross-species generalization of the model.

GANGE eliminates the need for high-coverage sequencing and parallel Illumina short-reads sequencing, something essential in all sequencing projects of today. Besides removing the errors, it also extends the read in both directions by 4kb with >92% average accuracy. Consequently, GANGE significantly reduces both the cost and time associated with genome sequencing projects while achieving highly reliable genome assemblies with better completeness. Its read extension capability has another very useful application: Decode promoter region in species whose genome is not sequenced, just by using their transcript sequence. When evaluated across 12 different species for promoter generation, including those not even represented in the training data, the system achieved accuracy levels >92%. This means that one can now also perform regulatory studies in such species for which genomic sequences are unavailable. The implications of GANGE extend far beyond the technical innovations. By reducing the reliance on costly hybrid sequencing workflows, it presents a transformative alternative to traditional genome sequencing and assembling pipelines, offering scalability and cost efficiency. Researchers working on non-model or underrepresented eukaryotic species, where reference genomes are incomplete or unavailable, can now benefit tremendously from this. GANGE also makes sequencers like ONT to become highly useful for large genome projects, achievable now at much lesser cost, opening the doorways for a huge revolution in the field of genome sequencing projects. **Figure 1** illustrates an abstract view of the introduction of the problem and the solution.

**Figure 1:**
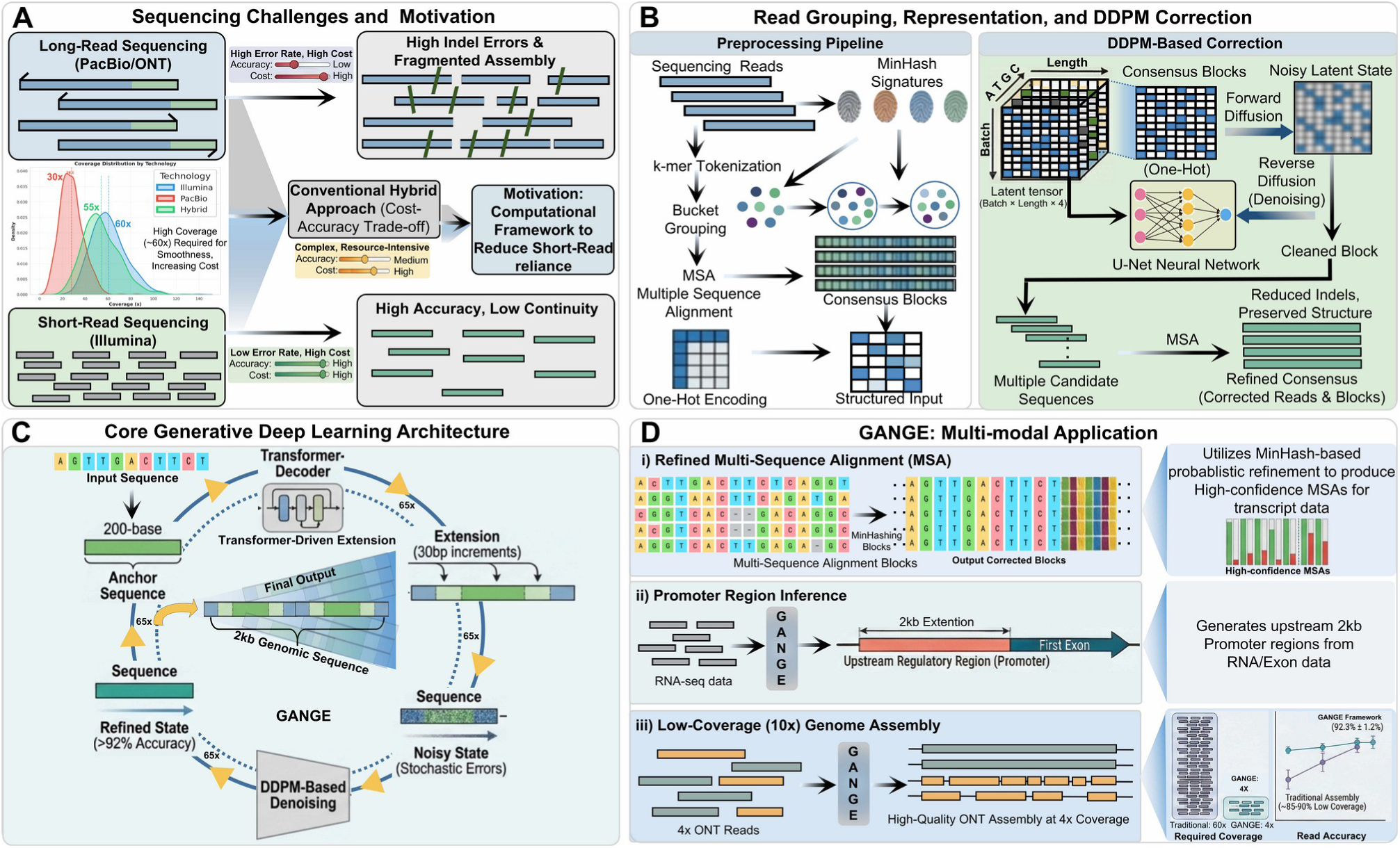
The workflow of the GANGE system. GANGE has four different levels.**(A)** The image illustrates the trade-off between the high continuity but high error rates of long-read sequencing (PacBio/ONT) and the high accuracy but low continuity of short-read sequencing (Illumina) by introducing a computational pipeline that reduces reliance on hybrid and high coverage sequencing. **(B)** First, the raw ONT reads are searched for any similarity overlap using ultra-fast Minhashing algorithm. This forms the clusters of shared similarity reads. In the next stage the clustered similarity shring reads undergo MSA. The raw MSA is refined for more correct ones using a novel k-meric iterative stepping realignment. The next stage takes this refined MSA to the DDPM guided error correction process where the noisy error prone positions are corrected by recovering the right bases. **(C)** The final stage is an iterative horizontal genration of upstream and downstream regions upto 2kb at each end. Based on the corrected consensus a dual Transformer-DDPM generative system generates the sequences through a 30 bases iterative cycles of generation and correction. **(D)** This way GANGE generate sequences with >92% accuracy from indel errors prone ONT reads at as low as 4X-10X depth and extends them further by 4kb. One can also use it to generate promoter sequences with same accuracy in species whose genome is unknown, just using RNA data.

## Materials and Methods

### Datasets construction

For the development of universal and generalized model for the error correction and extension of ONT reads, we retrieved genomes of 16 different eukarotic species i.e., *Oryza sativa, Zea mays, Arabidopsis thaliana, Vitis vinifera, Actinidia chinensis, Arabidopsis lyrata, Amborella trichopoda, Brassica napus, Citrus clementina, Cucumis sativus, Glycine max, Crocus sativus, Hordeum vulgare, Musa acuminata, Caenorhabditis elegans, and Drosophila melanogaster* from Ensembl Plants and NCBI **(Table S2)**. Other than genomic sequences, annotations, and gene sequences were downloaded from Ensembl Plants and NCBI. 2kb promoter sequences and their corresponding downstream genic sequences were extracted from gene transfer format (GTF) files for the collected species i.e., *O. sativa, Z. mays, A. thaliana, and V. vinifera* were finally integrated which has been named as Dataset “A” in this study. This dataset was used for the development of Transformer model. These species span monocots and dicots and include both model and non-model organisms, thereby capturing a wide range of genome sizes, repeat compositions, and regulatory architectures. The dataset having sequences from *A. chinensis, A. lyrata, A. trichopoda, B. napus, C. clementina, C. sativus, G. max, Crocus sativus, H. vulgare, M. acuminata, C. elegans, and D. melanogaster* have been named as Dataset “B” in the current study, used completely as the test set. Four species i.e., *O. sativa, Z. mays, A. thaliana, and V. vinifera* genomic sequences were taken and chopped into k-mers of 64 bases in non-overlapping manner for the creation of another dataset known as Dataset “C” in the current study. This dataset was used for the development of DDPM model. Dataset “A” and “C” were split in a 70:30 ratio for training and testing purposes, including for 10-folds random train:test trials. In the entire study this was ensured that no instance of training and testing overlapped in any manner, in order to ensure unbiased learning and performance.

### Biologically informed MinHash-based clustering of ONT reads

To organize noisy ONT reads in biologically cohesive manner before performing the deep-learning based error correction and assembly, we implemented a MinHash-based **[59]** read clustering strategy that grouped the ONT reads according to shared genomic origin. Adapter-trimmed and quality-filtered reads were transformed into overlapping *k*-mers, which encode local nucleotide composition and sequence context even when individual bases are affected by sequencing errors.

### Shingling and set representation

The first step in the MinHash pipeline was to convert each ONT adapter-trimmed and quality-filtered read from a linear nucleotide string into a set-based representation amenable to efficient similarity computation. This conversion was achieved by generating overlapping k-mers out of the reads.

Let, R = {r₁, r₂,…, rₙ} denote the complete set of reads, where each read ‘r_*i*_’ is a string over the nucleotide alphabet Σ = {A, C, G, T} of length ‘|r_*i*_|’ bases. For each read ‘r_*i*_’, a k-mer shingling function ϕₖ: Σ’ → 2^(Σ^k) was applied with k = 6 to extract the complete set of overlapping 6-mer subsequences. This function is formally defined as:

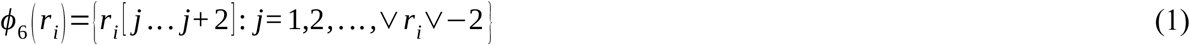

where, r_*i*_[j…j+2] denotes the substring of ‘r_*i*_’ beginning at position ‘j’ and spanning 6-mer. For a read of length ‘L’, this produces L - 5 overlapping 6-mers. These 6-mers were drawn from a possible vocabulary of |Σ|^6^ = 4^6^ = 4096 distinct 6-mer combinations.

Each read ‘r_*i*_’ was thus represented as a shingle set S*i* = ϕ₃(r_*i*_) ⊆ Σ³, and the full dataset was represented as the collection of shingle sets ℒ = {S₁, S₂,…, Sₙ}. The choice of k = 6 was motivated by a comprehensive analysis of error rate which is explained further in next sections.

### Jaccard similarity

Once each read had been converted into its shingle set representation, the pairwise sequence Jaccard similarity between any two reads could be computed using a set-theoretic metric. The pairwise sequence similarity between any two reads ‘r_*i*_’ and ‘r_ⱼ_’ was quantified using the Jaccard similarity **[60]** coefficient J(S_*i*_, S_ⱼ_), defined formally as the ratio of the cardinality of the intersection to the cardinality of the union of their respective shingle sets:

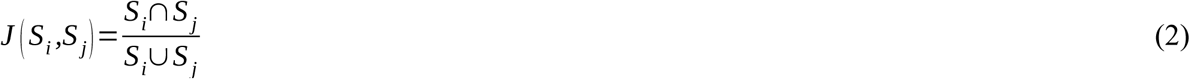

where J(S*i*, Sⱼ) ∈ [0, 1], with J = 0 indicating complete dissimilarity the two reads share no 6-mers whatsoever and J = 1 indicating identical shingle sets, implying that both reads contain exactly the same set of hexanucleotide subsequences.

It is important to note that direct computation of J(S_*i*_, S_ⱼ_) or all C(n, 2) read pairs required explicit construction and comparison of shingle sets, which scaled to O(n²L) in both time and space for ‘n’ reads of mean length ‘L’. For a typical ONT dataset containing hundreds of thousands of reads, this brute-force computation is computationally intractable. The MinHash algorithm described in the following section resolves this scalability problem by replacing exact Jaccard computation with an efficient random hash functions.

### MinHash signature generation

Given a uniformly random permutation ‘π’ of the element space Σ^6^, the probability that the minimum-valued element under ‘π’ is the same for two sets ‘S_*i*_’ and ‘S_ⱼ_’ is exactly equal to their Jaccard similarity. This can be stated formally as:

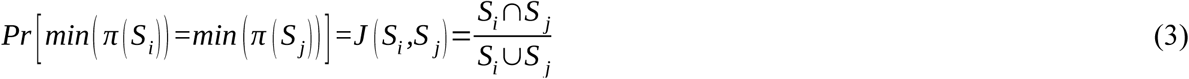

where, π(S) denotes the image of the set S under the permutation π, and min(π(S)) denotes the element of ‘S’ that maps to the smallest value under ‘π’. In practice, truly random permutations of the full k-mer space are computationally expensive to generate and store. Instead, they were approximated using a family of m independent hash functions H = {h₁, h₂,…, hₘ}, where each hash function hₜ: Σ³ → ℤ maps trinucleotide strings to integer values. For each read ‘r_*i*_’ and each hash function ‘h_ₜ_’, the MinHash signature value was computed as the minimum hash value attained across all 6-mers in the shingle set ‘S_*i*_’:

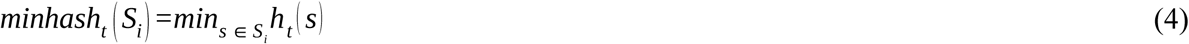

This operation identified the 6-mer in ‘S_*i*_’ that hashed to the smallest integer value under ‘h_ₜ_’, serving as a compact representative of the entire shingle set under that hash function. The complete MinHash signature of read ‘r_*i*_’ was now the vector of ‘m’ such minimum hash values, one per hash function:

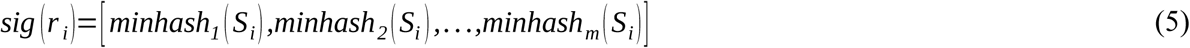

Thus, each signature here was therefore a fixed-length vector of ‘m’ integers regardless of the original read length ‘L’, compressing the variable-length shingle set into a compact, constant-size representation suitable for efficient storage and comparison. In this study, m = 128 hash functions were used, producing 128-dimensional signature vectors for each read. The hash functions were implemented as pairwise-independent linear hash functions of the form h_ₜ_(x) = (a_ₜ_ · x + b_ₜ_) mod ‘p’, where ‘a_ₜ_’ and ‘b_ₜ_’ are randomly sampled coefficients and ‘p’ is a large prime, ensuring the independence of hash function outputs required for accurate Jaccard estimation.

### Jaccard similarity estimation from signatures

Given the MinHash signatures of two reads ‘r_*i*_’ and ‘r_j_’, the Jaccard similarity J(S_*i*_, S_ⱼ_) can be estimated without ever explicitly constructing or comparing their full shingle sets. The estimated Jaccard similarity is defined as:

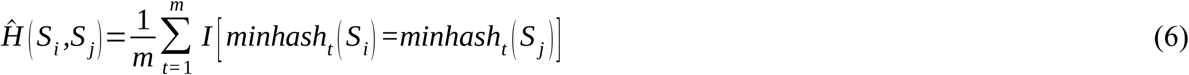

where, *‘I’* is a binary indicator function. The variance of this estimator decreases as ‘m’ increases, with the standard error of the estimate given by:

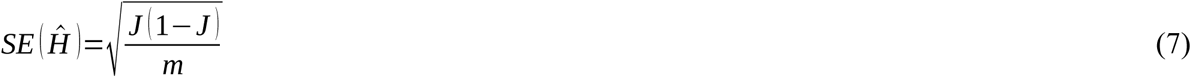

At m = 128 hash functions and a true Jaccard similarity of J = 0.7, the standard error was approximately √(0.7 × 0.3 / 128) ≈ 0.040. The computational complexity of similarity estimation was thereby reduced from O(|S*i*| + |Sⱼ|) for exact Jaccard computation to O(m) for signature comparison, where m = 128 is a small constant independent of read length.

Importantly, this alignment-free similarity measure tolerates the insertion-deletion-dominated error profile of ONT data, reads were clustered using a Jaccard similarity threshold of 0.7. These biologically coherent read clusters formed the foundation blocks effective coverage, accurate consensus generation, and error correction without requiring excessive sequencing depths as will transpire below.

### Refined multiple sequence alignments across the generated clusters

The clustered sequences were subjected to multiple sequence alignment (MSA) while using MUSCLE **[61]**. Let *Seq* = {s_1_, s_2_,…….., s_n_} be a set of clustered sequences and ‘A_0_’ be the initial alignment produced by the MUSCLE aligner. The motivation for the 6-mer refinement is based on the chance of observing the next indel after a given interval, where an error at position ‘j’ sets the mark for observing the interval after which observing the next indel was most likely. It was for *k*=6:

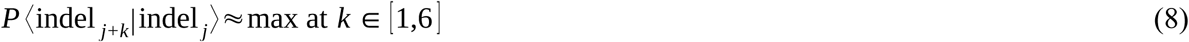

The decision of 6-mer was chosen after observing that probability of seeing an indel error after the previous event was highest around 6 bases (**Figure 2b**). **Figure 3** illustrates the MSA improvement methodology. The staged MSA refinement is defined by an iterative function where, in each stage ‘s’, a 6-mer block is truncated from the sequence and the remaining set is realigned. If *Seq* is the set of sequences, the transformation for each iteration is:

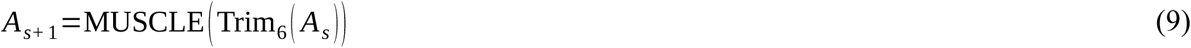

**Figure 2:**
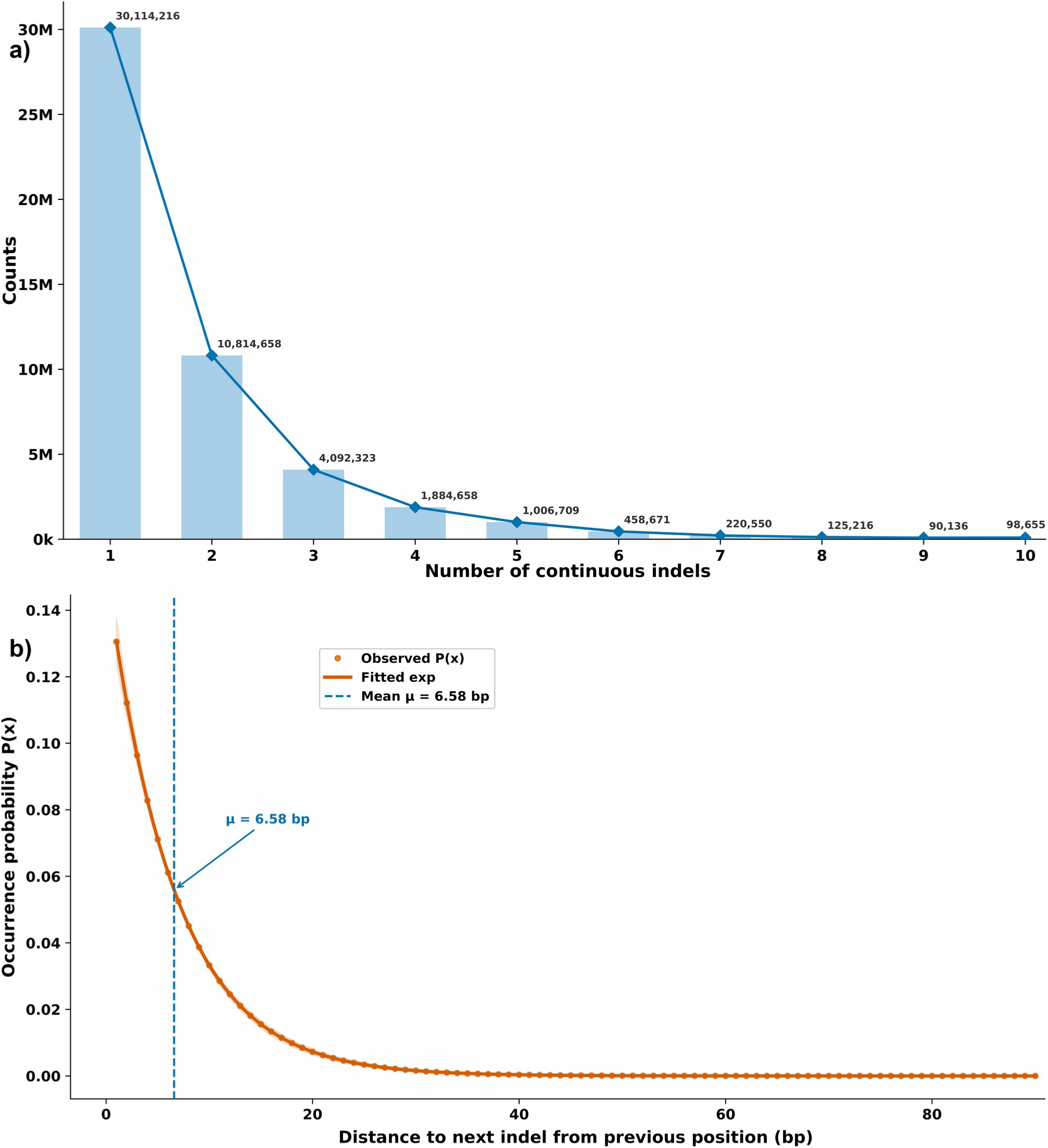
Distribution and probabilistic modeling of indels lengths in long-read reads. **(a)** Frequency distribution of indels lengths (1-10 bp) observed in long-read sequencing data. The counts decrease sharply with increasing deletion size, with single-base deletions being the most prevalent (∼30.1 million events), followed by 2-bp (∼10.8 million) and 3-bp (∼4.09 million) deletions. Indels longer than 5 bp occur substantially less frequently (<1 million events), demonstrating a strongly right-skewed distribution characteristic of sequencing-induced indel errors. **(b)** The probability distribution plot of seeing the next indel event. It emerged that the ∼6 bases interval is expected to see the next indel event. This helped to determine the suitable k-mer (6-mer) in which the results and MSA had to be broken and processed for MinHash based similarity search and MSA refinements.

**Figure 3:**
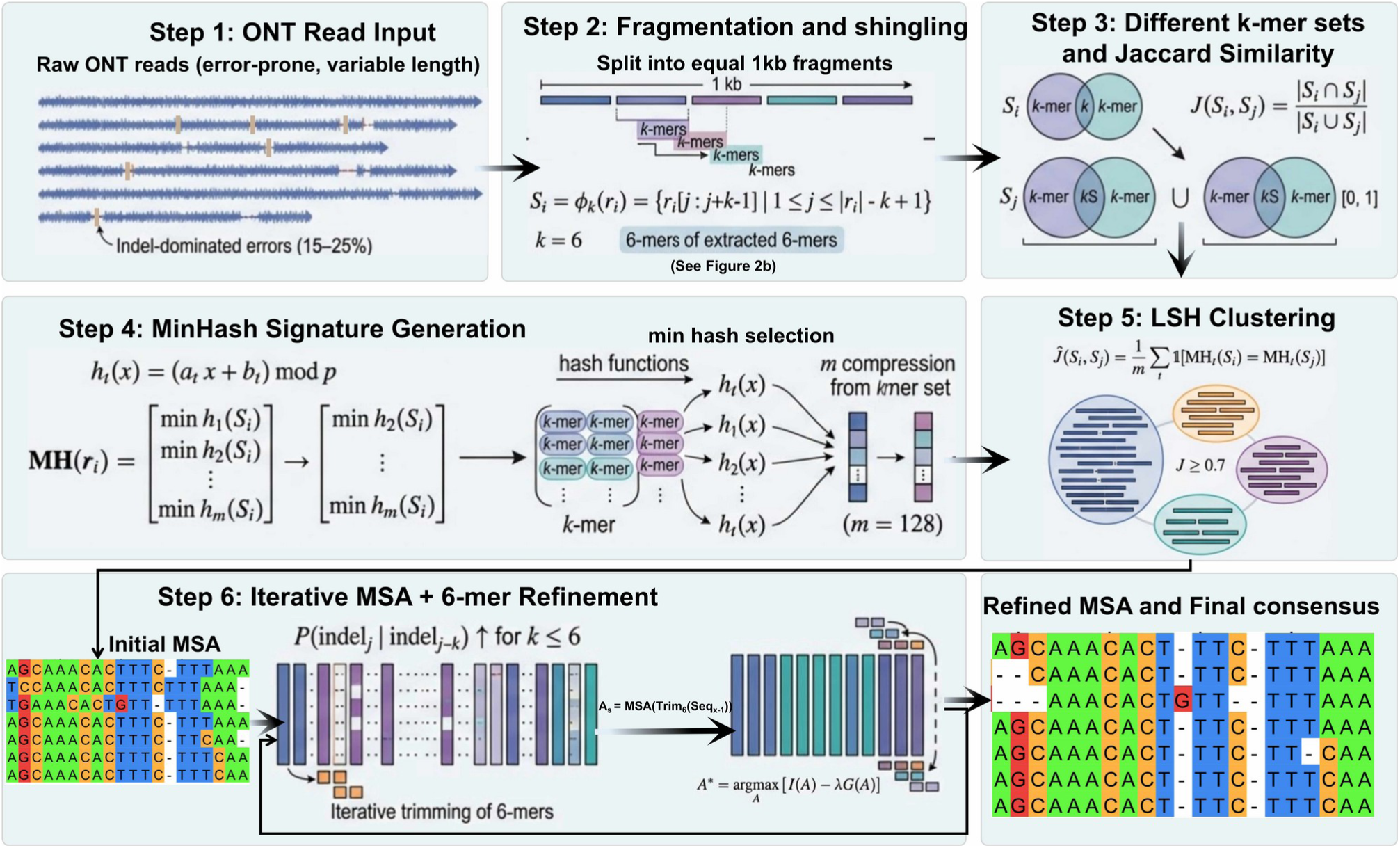
Workflow for MinHash-based clustering and iterative MSA refinement of long reads. The pipeline begins with the input of raw reads characterized by variable lengths and high indel-dominated error rates (Step 1), which are subsequently fragmented into equal 1kb segments and processed into 6-mer “shingles” (Step 2). To evaluate sequence relationships, Jaccard Similarity was calculated across different k-mer sets (Step 3). These sets are compressed into MinHash signatures using m=128 hash functions to identify minimum hash values for each sequence (Step 4), allowing for efficient Locality-Sensitive Hashing (LSH) clustering of reads with high similarity (Step 5). Finally, an iterative Multiple Sequence Alignment (MSA) and 6-mer refinement process is applied leveraging the probability of indels to trim error-prone k-mers to produce a refined MSA and a high-accuracy final consensus sequence (Step 6).

This iterative “shaving” of the sequence ends ensures that edge-case alignment artifacts, which often anchor indel errors incorrectly, are recalculated. The optimal alignment ‘À’ is reached when the biological information ‘I(A)’ is maximized relative to the gap penalty ‘G’:

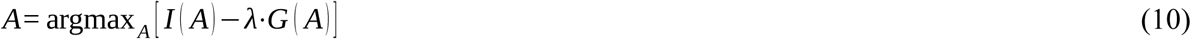

The refined MSA creates a structured grid where aligned reads ‘R’ and error spots ‘E’ are positioned relative to a valid consensus ‘C’. This corrected matrix serves as the high-fidelity training data for the model to learn the transition from an “errored spot” to a “valid base” based on the local alignment context. These above steps ensured a far better much improved and biologically more informative MSA, while countering the indels errors influence. A correct MSA ensured blocks of aligned reads and aligned error spot placed in validity for more correct consensus formation, which in otherwise missed by general MSAs. Despite all of this, certain positions within the MSA inevitably exhibited a distribution of nucleotides that was effectively random that is, where no single nucleotide achieved statistically dominant representation. Such positional profiles are the molecular signature of genuine uncertainty, arising most commonly from the errors present in ONT reads besides clear indels. The resulting one-hot or soft-encoded positional vector carries no reliable discriminative signal and instead encodes pure uncertainty a state informationally equivalent to noise within the tensor representation. Such positions were therefore explicitly designated as noise sites requiring correction by the DDPM. This designation is biologically meaningful: rather than propagating false high-confidence base assignments at positions where the sequencing and alignment evidence is fundamentally ambiguous, explicitly labeling these positions as uncertain allows the denoising diffusion process to apply its learned prior over nucleotide distributions to recover the most probable true base state, guided by the broader sequence context. After all this, the corrected MSA forms the data-points to learn from and generate correct bases at the errored spots.

### Representation of corrected MSA blocks for DDPM

A MSA block generated after the corrective measures became like an image file where consistent blocks were like pixels (data points) where no error/missing data was existent. While the positions having indels or inconsistent base allocation became the spots/pixels where the suitable base had to be recovered through generation done by DDPM **[62]**.

To enable diffusion-based modeling, each aligned block was transformed into a structured numerical representation. Each nucleotide position of a sequences were encoded using the representation where A = [1, 0, 0, 0], C = [0, 1, 0, 0], G = [0, 0, 1, 0], and T = [0, 0, 0, 1], while ambiguous bases or gaps such as “N” or “-” were assigned an all-zero vector to denote uncertainty to reflect equal probability across nucleotides. Aligned blocks were partitioned or padded into fixed-size windows and reshaped into three-dimensional tensors of size 64 × 64 × 4, providing a uniform input format for stable DDPM training. A row-major mapping strategy was used to fill sequences left-to-right and top-to-bottom within the grid, ensuring that local nucleotide adjacency and alignment context were preserved in the spatial layout. Shorter sequences were zero-padded, while longer alignments were segmented into overlapping windows to ensure complete genomic coverage.

In case, where the number of contributing sequences for MSA blocks were below 64, such cases were flagged and handled through a dedicated low-coverage protocol. In such cases, where the number of aligned sequences were critically low, the consensus signal becomes statistically unreliable, as individual sequencing errors cannot be distinguished from true biological variation without sufficient read redundancy. To mitigate the statistical unreliability introduced by insufficient read depth, a coverage augmentation strategy was employed **[63–64]**. The duplicated sequences were subjected to controlled stochastic perturbation introducing low-magnitude random noise into the one-hot encoded vectors prior to tensor construction. This augmented tensor was then processed through the standard 64 × 64 × 4 encoding pipeline identically to well-covered alignment blocks, ensuring architectural consistency across all input batches. This stratified handling of variable-depth MSA regions ensured that the DDPM received consistent, information-rich input representations across the full range of sequencing depths encountered in real long-read datasets, while explicitly preventing low-coverage alignment blocks from corrupting the learned denoising distribution.

Within the DDPM framework, these sequence blocks were treated as structured biological signals undergoing controlled corruption through a forward diffusion process. Noise was progressively added to nucleotide states while preserving positional alignment, mimicking the stochastic error processes observed in long-read sequencing. During reverse diffusion, the model learns to reconstruct the original aligned block by leveraging contextual information across neighboring positions and across homologous reads within the cluster. Because the input blocks originate from biologically coherent MSAs, the DDPM implicitly learns the conservation patterns, local sequence grammar, and error signatures simultaneously.

### Diffusion model framework to reclaiming the errored sequences

The above steps ensured correctly aligned reads and correctly positioned errors, all in a block form. This block may be seen as an image where the errored positions were the spots for recovery of correct bases. And this is what exactly DDPM achieves for corrupted or missing data: Generate the most probable data-point sample, based on a proposal density. DDPM operates in two phases: **i)** Step wise corruption on input data by adding Gaussian noise in stochastic manner, **ii)** Learning the noise, and recovering back the original data from the corrupted noisy input.

### The Denoising Diffusion Probabilistic Model for sequencing reads correction

The corrected aligned blocks were segmented into fixed-length patches of sequence length *L* = 64 bases, leveraging the 10X coverage to stack multiple overlapping reads per position.

Each base in the sequence is represented via one-hot encoding (OHE) over the alphabet [***A***, ***T***, ***G***, ***C***] yielding a 4-dimensional channel for nucleotide identity, treating the data as a multi-channel “image” analogous to pixel intensities in image generation tasks. This transforms the input into a tensor *X* ∊ ℝ^64×64×4^.

In matrix form, ‘**X_0_*’*** can be expressed as a block-structured tensor:

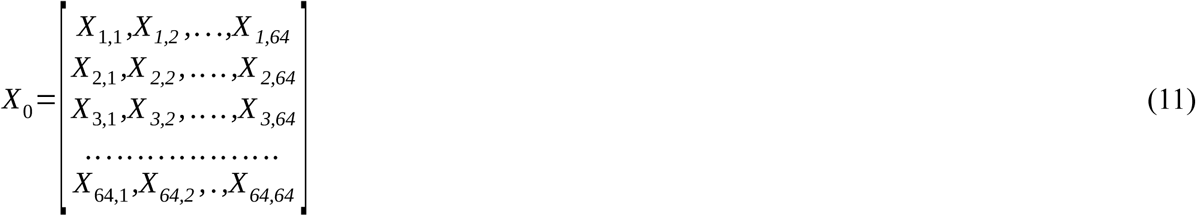

where each *X* ∊ ℝ^4^ is the OHE base vector at position ‘i’ under coverage stack ‘j’. This representation preserves local sequence context while enabling convolutional processing in the DDPM, akin to treating sequencing errors as pixel-level noise in an image.

For conditional error correction, the DDPM is conditioned on a reference sequence patch *c*∊ ℝ^64×1×4^ (OHE aligned), injected via cross-attention in the U-Net backbone during both training and inference. The U-Net backbone processes this conditional input through a symmetric hierarchy of residual and attention blocks, utilizing a [1, 2, 2, 4] channel multiplier to scale feature depth from 128 to 512 channels.

### Forward diffusion process

The forward diffusion process models the gradual corruption of the latent representation ‘z_0_**’** of the correct MSA blocks at the start of the diffusion process into pure noise over ‘T’ = 1000 discrete time steps, interpreting ONT sequencing errors as initial “noise” that is progressively amplified.

This is a fixed Markov chain 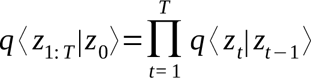, where each transition adds isotropic Gaussian noise:

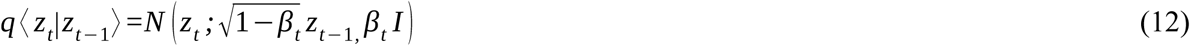

with *β*∊ ( 0, 1) as the variance schedule (detailed below). The chain destroys structure such that *z_t_* ≈*N* (0, 1), simulating the transition from a corrected sequence to a fully erroneous (noisy) observation. ‘z_t_’ represents the progressively noisier latent state of the MSA blocks at a specific diffusion timestep ‘t’, resulting from the controlled addition of Gaussian noise.

A closed-form expression allows direct sampling from *q* ⟨ *z_t_*|*z*_0_ ⟩ without sequential computation, leveraging the reparameterization trick:

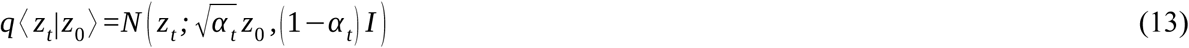

where, 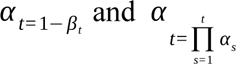. Reparameterizing with standard Gaussian noise *ε*≈ *N* (0 *, I*):

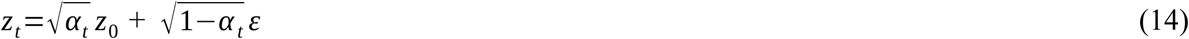

In matrix form, for a flattened latent *z*_0_ ∊ ℝ (where *N=d* ×4), this is:

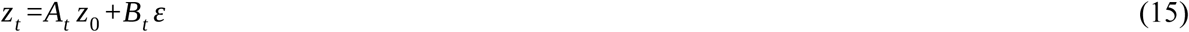

with 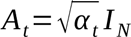 (diagonal scaling matrix) and 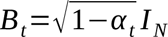. This enables efficient batch training by sampling *t* ∼Uniform {1,…*, T* } and *ε* per minibatch.

### Variance scheduling

The noise schedule 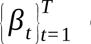 controls the rate of corruption, ensuring gradual signal degradation over *T=*1000 steps. A linear schedule was used:

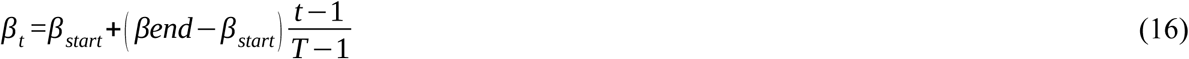

with β*_start_*=10^−4^ and β*_end_*=0.02, yielding small increments early (preserving structure) and larger later (accelerating to isotropy). Precomputed cumulatives include:

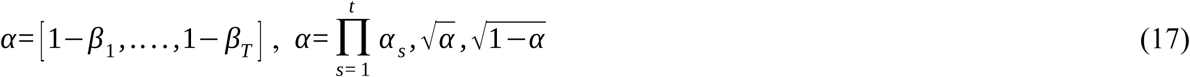

The posterior variance for the true reverse transition *q* ⟨ *z_t_* _−1_|*z_t,_ z*_0_⟩ is:

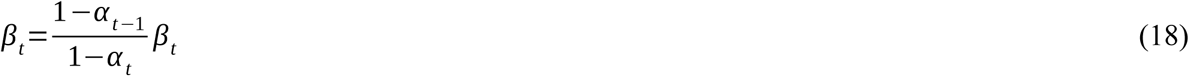

used to parameterize the learned reverse variance (see below).

In the ONT context, this forward process augments real sequencing errors with synthetic Gaussian noise, training the model to “hallucinate” corrections by inverting the corruption.

### Reverse diffusion process

The reverse process learns to denoise from *z_t_* ≈*N* (0 *, I*), back to ‘z_0_***’***, parameterized as a conditional Markov chain 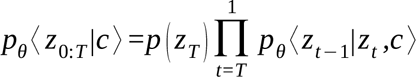, where conditioning on the reference ‘c’ guides correction toward the true sequence. Each step is a Gaussian:

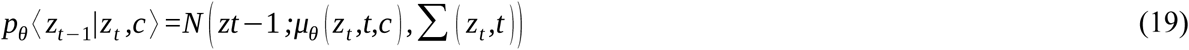

with fixed isotropic variance _∑_(*t*)*=β_t_*. The mean is derived from noise prediction *ε_θ_* _(_ *z_t_, t, c* _)_≈*ε*:

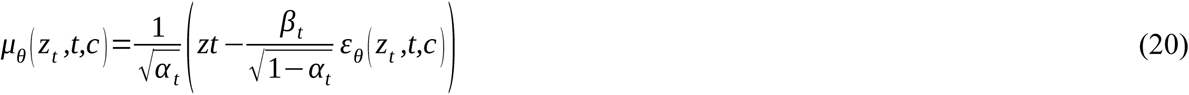

Equivalently, predict the clean latent:

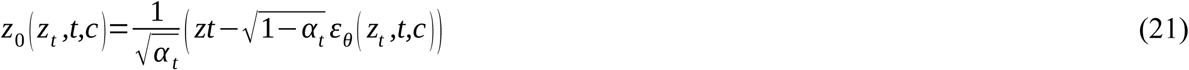

then, 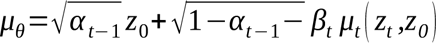, where *μ_t_* is the posterior mean.

### Training

The model *ε_θ_* (a U-Net with time embedding and cross-attention to ‘***c’***) is trained via simplified variational lower bound, reducing to noise prediction MSE, with *t*∼*Uniform* {1,…, 1000 }. This leverages the reparameterization for low-variance gradients. In matrix form, for batched inputs ‘ Z_t_’ ∊ ℝ*^B^*^×*N*^ (batch size B).

The U-Net architecture mirrors standard DDPM implementations: time ‘***t’*** is sinusoidally embedded and projected via MLP, fed into residual blocks with GroupNorm and Swish activations. Down/upsampling used 4 levels (channels: 64→128→256→512), with self-attention at intermediate resolutions and cross-attention to ‘c’ (projected to keys/values). Output: Conv2D to 4 channels predicting *ε_θ_*.

### Sampling for error correction

Inference starts from the noisy latent *z_T_* ≈*N* (0 *, I*) with iterative denoising over ***T*** = 1000:

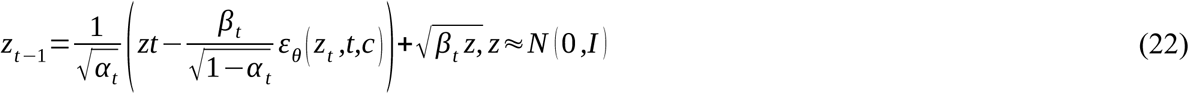

with *z=* 0 *, t=*1. The final ‘***z_0_’*** is decoded to corrected *X*_0_ *=argmaxp_θ_* ⟨ *X*_0_|*z*_0_ ⟩ (via softmax over 4 channels). For ultra-long ONT reads, patches were processed sequentially with overlap, aggregating via consensus.

This conditional DDPM framework achieves error correction by learning to reverse sequencing noise, with the 1000-step schedule ensuring fine-grained refinement and the latent space enabling tractable computation for 64×64×4 inputs.

### Evaluation of the generated output from the DDPM reverse process

The DDPM reverse process generates multiple sequences representing corrected reads, which are then used to draw the consensus sequence which is matched to the original target sequences. Consensus sequences are derived from each block using majority voting of base calling, producing corrected representative sequences. Post-decoding, the model’s accuracy is evaluated by comparing the generated sequences to actual sequence, analyzing base-by-base agreement.

The generative process is defined by the reverse transition probability ‘p_θ_’, which reconstructs a cleaner sequence ‘x_t-1_’ from a noisier state ‘x_t_’ through learned Gaussian parameters: The reverse process transforms a noisy latent representation ‘x_T_’ back into a clean sequence ‘x_0_’ through a series of learned transitions:

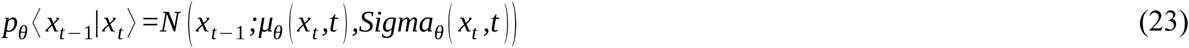

By iteratively applying this process starting from pure noise ‘x_T_’, the model generates a set of ‘N’ candidate sequences, ***S*** = {s_1_, s_2_, s_n_}.

For each genomic block, a consensus base ‘B’ at position ‘j’ is derived by selecting the most frequent character across the ensemble of ‘N’ generated sequences. This is expressed using the argmax of the indicator function sum:

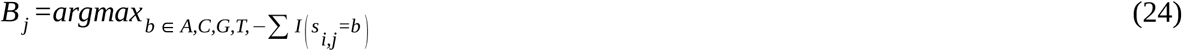

Where, ‘b’ is the set of possible bases and the gap token. I(.) is the indicator function, which equals 1 if the condition is true and 0 otherwise. The performance of the post-decoding phase is quantified by calculating the normalized agreement between the DDPM-derived consensus sequence ‘S_cons_’ and the experimental target sequence ‘S_target_’:

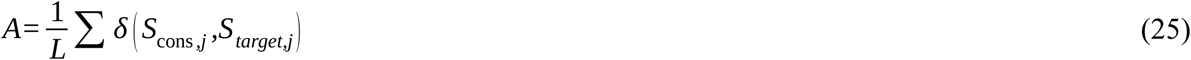

where, ***L*** is the total sequence length. δ(a, b) is the Kronecker delta function, returning 1 if a = b else 0.

This entire part of the study has used 70% of the Dataset “C” as the training set to build the model and its performance was first reassured across the remaining 30% totally unseen test dataset instances. **Figure 4** depicts the operation of the implemented deep-learning DDPM system. Further to this, the evaluation was also done through 10-fold random independent train-test trials while building the model from the scratch every time maintaining the same 70:30 ratio of train:test datasets to evaluate the consistency of the developed method across variable training and testing datasets.

**Figure 4:**
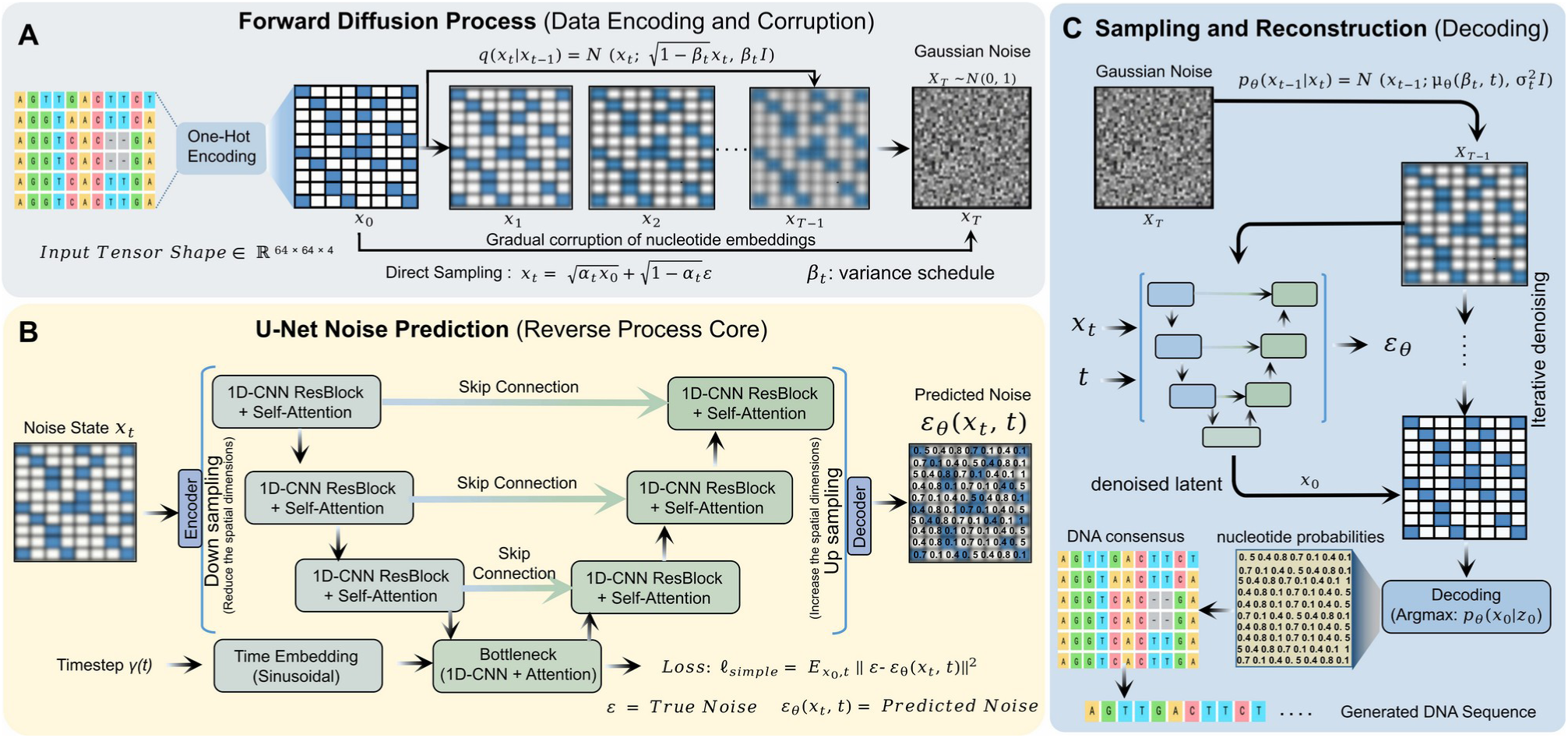
The DDPM architecture. The DDPM architecture is divided into three core stages: **(A)** Forward Diffusion Process, where input DNA sequences are one-hot encoded into a 64X64X4 tensor and systematically corrupted with Gaussian noise over T timesteps according to a variance schedule, transitioning from a clean state to pure noise. **(B)** U-Net Noise generation, which serves as the reverse process core, utilizing an encoder-decoder structure with 1D-CNN ResBlocks and self-attention mechanisms to generate the noise added at any given timestep, guided by sinusoidal time embeddings and optimized via a loss function, and **(C)** Sampling and Reconstruction, where the model iteratively denoises a random Gaussian noise starting point to recover the denoised latent, which is then decoded using an Argmax function on the generated nucleotide probabilities to generate a high-fidelity DNA consensus sequence.

### The Transformer system

Dataset “A” encompassed the extracted 2kb promoter sequences covering 135, 858 genes from 4 species (*O. sativa, Z. mays, A. thaliana, and V. vinifera*) to develop transformer system. This part enables the model to take sequences data as input and capture both local and global patterns in the DNA sequences for generation of 2kb upstream sequences. The transformer architecture was implemented using PyTorch. The reason for selecting promoter regions was that these 2kb upstream sequences host most of the regulatory components governing the expression of the downstream genic region. Thus, using this transformer, one can conduct regulatory studies even in species with unsequenced genomes.

### Word representations of sequence data for Transformer

The DNA alphabet consists of four nucleotides: A, T, G, and C, forming a total of 16 unique dimers. Similarly, this alphabet can create 1, 024 unique 5-mer words, 4, 096 unique 6-mer words, and 16, 384 unique 7-mer words. In processing these sequences, overlapping windows of pentamers, hexamers, and heptamers can capture critical information about structural and functional regions within the DNA **[65–66]**. The Transformer takes two inputs: source and target input. The source input consists of sequences with a maximum length of 585 words spanning 200 bases. The target input was their corresponding promoter sequence regions. This input had a maximum length of 24 words and spanned 30 bases. Each unique word was tokenized with unique numerical value, later converted into numeric vectors (embedding). The encoder processes these vectors to understand the input while the decoder uses that understanding to generate the output step-by-step. By feeding these embedded sequences into the model, the Transformer learned to translate the sequences in the form of generated upstream sequences **[67]**.

### Implementation of the Transformer’s Encoders-Decoders

The encoder reads and deciphers the grammar hidden in input sequence. It does so in layers which analyze how all the words relate to each other. Thereafter, the decoder uses this learning to build the output sequence word-by-word. It uses the context derived from the encoder’s outputs and incorporates its own attention mechanisms to focus on the relevant parts of the input sequence while generating each token of the output.

### The Encoderv

The embedding layer projects discrete input word tokens into continuous vector embeddings. The embedding layer is represented as a matrix *E_embed_* ∊ ℝ*^v^*^×*d*^.

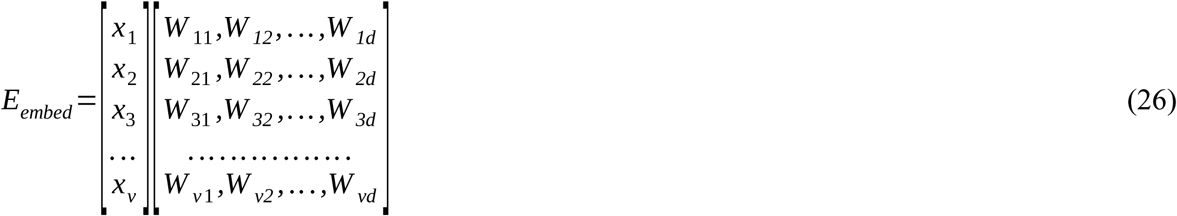

where, ‘v’ = vocabulary size and ‘d’ = embedding dimension.

Each word in matrix “*E_embed_”* combines with its positional embedding “*P”*, where “*P”* shares dimension “*d”* with the word embedding vector. The resulting matrix, “*E’_embed_”*, is obtained through “ *E’_embed_ =E_embed_ +P*”, where “*P”* is calculated using sinusoidal positional embedding equations:

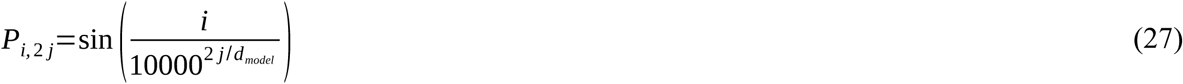

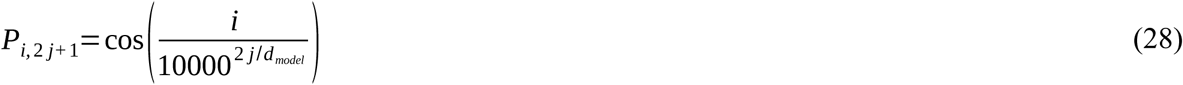

Here, “*i*” represents the position of the token, and “*d_model_*” is the embedding dimension. “*E’_embed_*” matrix enters the transformer encoder block, which processes it through a multi-head attention layer. The multi-head attention layer result is obtained by concatenating ‘*h’* individual attention heads and (head_1_, head_2_, head_n_) multiplying them by a weight matrix ‘*W^0^’*.

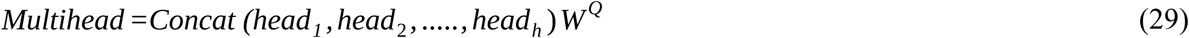

Each attention head (*head_i_*) calculates the attention mechanism with input matrices ‘Q’, ‘K’, and ‘V’. These input matrices are derived from “*M’”* using optimizable weight matrices ‘*W^Q^’*, ‘*W^K^’*, and ‘*W^V^’*.

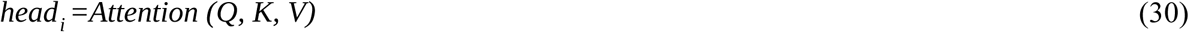

where, Q = M’.*W^Q^*, K = M’.*W^K^*, V = M’.*W^V.^.* The attention operation itself is defined by:

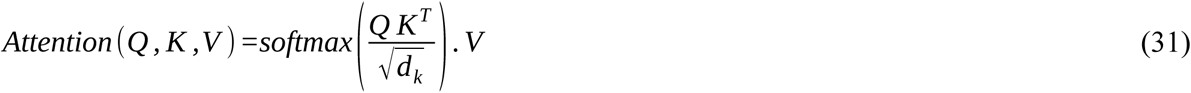

This entire process is applied to input matrices ‘Q’, ‘K’, and ‘V’. Finally, the ‘ h’ attention head outputs are concatenated and further transformed using an output weight matrix ‘W^0^’. The concatenated attention vectors are forwarded to a feed-forward network block, dropout layers, and a normalization layer. The output post-normalization is again directed to fully connected feed-forward layers, a dropout layer, and another normalization layer. The final output from this enhanced encoder structure feeds into the decoder block of the transformer.

### The Decoder

The decoder builds the output sequence one word at a time, using three main parts. First, the masked multi-head self-attention mechanism is essential for ensuring that the likelihood for a particular token depends only on the tokens that have been generated so far, thus preventing any information leakage from future tokens.

### Masked multi-head attention of decoders

For each distinctive head “*i*”, symbolizing *i=* 1, 2,…, ℎ, the calculation of three pivotal matrices ensues: “*Q_i_*” (Query), “*K_i_*” (Key), and “*V_i_*” (Value). This is accomplished through the matrix multiplication of “*E*” with the corresponding weight matrices “*W_qi_*”, “*W_ki_*”, and “*W_vi_*”. The formulation of attention scores “*Z_i_*” transpires through the application of the softmax function to the scaled dot-product attention mechanism, adhering to the expression:

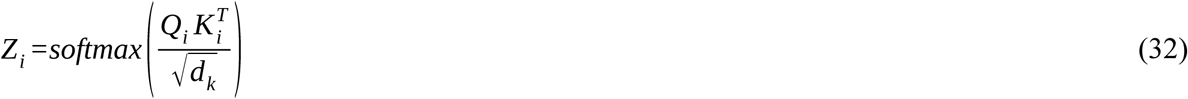

To preserve causality during training, a masking mechanism is applied to the attention scores, specifically designed to impede the model from attending to future positions, encapsulating the essence of masked self-attention. The ensuing phase involves the computation of the weighted sum, where each head “*i*” contributes to the formation of “*Head_i_*” via the formula:

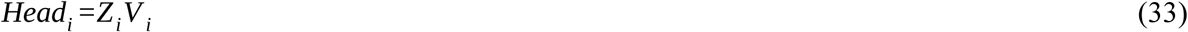

A concatenation of the outputs from all heads happens, culminating in a subsequent multiplication with the output weight matrix “*W_o_*”, yielding the definitive output of the Masked Multi-Head Attention mechanism in the decoder:

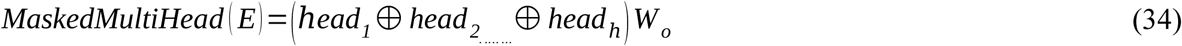

This exposition details the steps involved in using masked self-attention across multiple heads within the decoder of a Transformer model. During the decoding phase, the process begins with the Masked Multi-Head Attention mechanism, which analyzes the input sequence while ensuring that future tokens remain concealed. Once this initial processing is complete, the output from the encoder is integrated as a secondary input for the decoder. Next, the Decoder Multi-Head Attention mechanism operates similarly to the encoder’s attention, allowing the model to selectively focus on specific elements of both the input sequence and the encoder’s output. Afterwards, the output is refined through two Feed-forward and one normalization layer. Building on the normalized output, the model calculates a conditional probability distribution over the vocabulary for the next token in the sequence, using the softmax function. The resulting probabilities guide the selection of the next token. The conditional probability *P* ⟨ *y_t_*|*y*_1_,….*, y_t_* _−_*_1_, x* ⟩ for the next token “*y_t_*” is computed using the following equation:

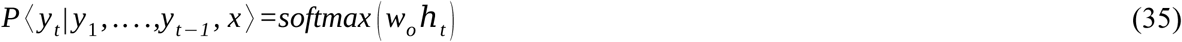

where, “*w_o_*” is the weight matrix, and “*h_t_*” is the hidden state at position “*t*”. This process repeats iteratively for each time step until the entire sequence is decoded. The model uses the generated tokens as input for subsequent time steps, allowing for autoregressive decoding.

Developed by Papineni et al, in 2002 **[68]**, Bilingual Evaluation Understudy (BLEU) score quantifies how closely a machine-generated sequences output aligns with target sequences. The final BLEU score is computed as the weighted geometric mean of the precision scores of 5-grams levels in the generated sequence compared to the reference sequence, adjusted by the brevity penalty. Mathematically, this is represented as:

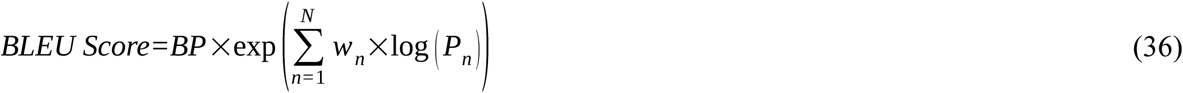

where, “*BP*” is the Brevity Penalty, accounting for the length of the generated sequence relative to the reference. “*N*” is the maximum order of 5-grams considered. “*w_n_*” is the weight assigned to each n-gram precision. “*P_n_*” is the precision of 5-grams in the predicted sequence. The primary advantage of the BLEU score is its ability to provide a quick, quantitative assessment of translation quality. Its reliance on n-gram matching makes it easy to compute and interpret. **Figure 5** depicts the operation of the implemented deep-learning encoder-decoder system.

**Figure 5:**
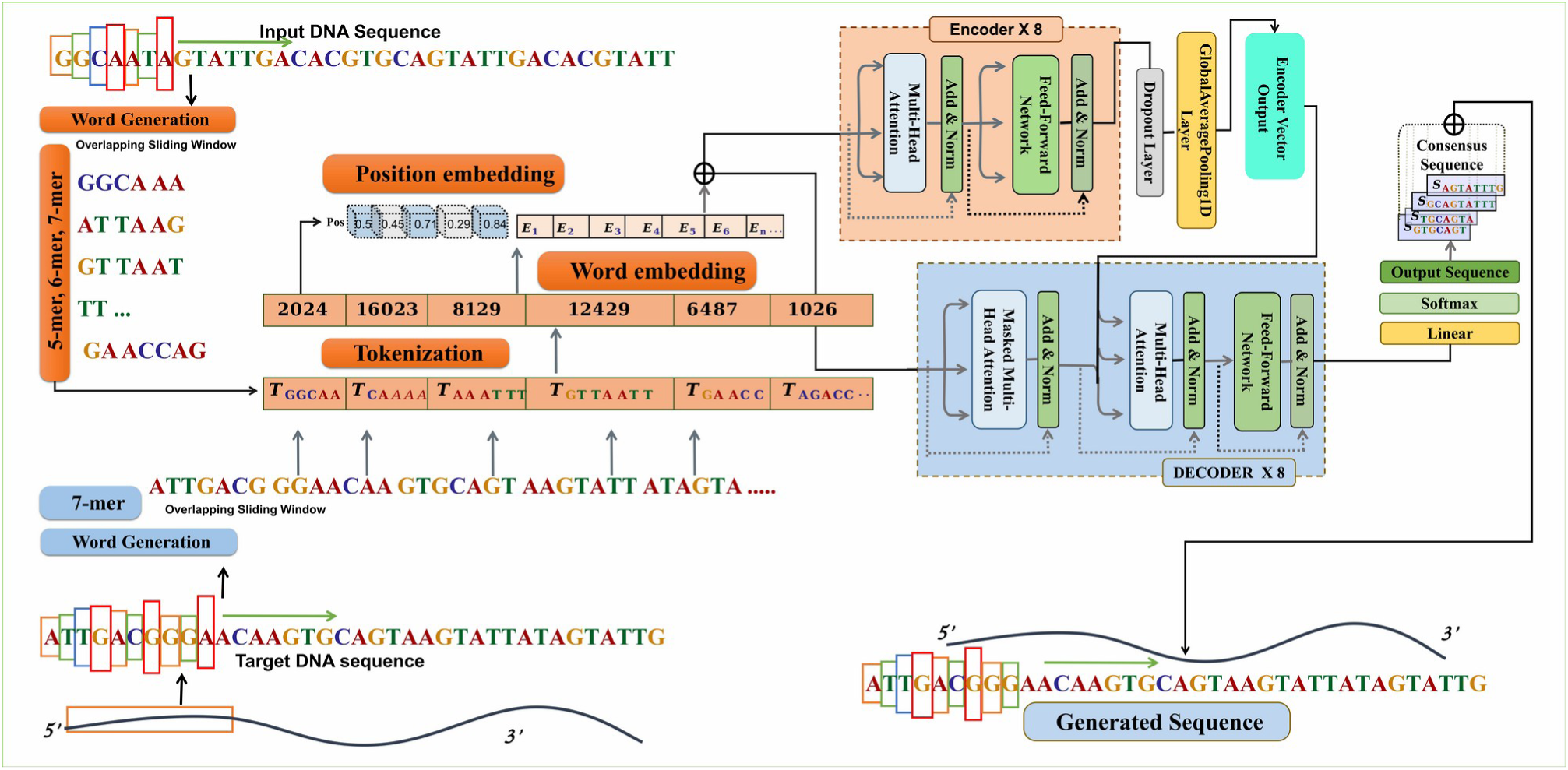
Implementation of the Transformer system to generate promoter sequence. The encoder processes sequences represented in pentamer, hexamer, and heptamer word representations, and the decoder receives the upstream sequence information for the corresponding sequence. Eight layers of encoder–decoder pairs are incorporated. The learned representations from the encoder were passed to the encoder–decoder multi-head attention layer, which also receives input from decoder layers. The resulting output then undergoes conditional probability estimation, which plays a pivotal role in the decoding process. After layer normalization, the model proceeds to calculate the conditional probability distribution over the vocabulary for the next token backed by a attention mechanism. Finally, the resulting tokens are converted back into words, representing the promoter sequence for the given sequence.

### Recursive horizontal sequence generation by Transformer

The generative capacity of the Transformer-Decoder was evaluated based on its ability to perform iterative sequence extension, transforming short-range genomic inputs into long-range contiguous blocks. In this framework, the model generates a 30-base extension ‘δS’ from an input window ‘W’ of 200 bases. This is defined by the conditional probability of the sequence ‘S’ at step ‘i’:

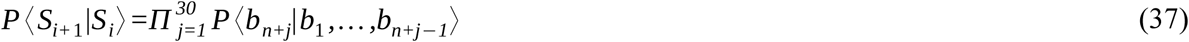

Where, ‘b’ represents individual nucleotides and ‘n’ is the current sequence length. For the full 2kb flanking region, the recursive stitching strategy follows the transition:

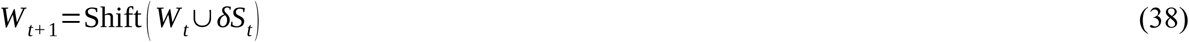

Within a single inference cycle, the likelihood of a generated 30-mer is synergistically derived from the overlapping *k*-mer distributions (***k ɛ {5,…., 7}***). The total log-likelihood ‘L’ for a generated segment is the weighted sum of these grammars:

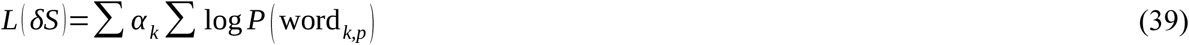

Here, ***α_k_*** represents the importance weight assigned to each *k*-mer scale, ensuring that both local and complex *k*-mers are preserved. The final accuracy of the 2kb promoter sequence ‘S_gen_’ is calculated by a per-base comparison against the experimental ground truth ‘S_exp_’ using an indicator function:

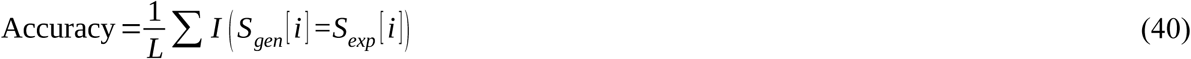

Where, ***L*** = 2000 and ***I*** is 1 if the nucleotides match, else 0. This provides a granular validation of the model’s structural integrity across the entire generated block.

In this way, the accuracy of the raised model was calculated for the complete promoter output sequence. We had previously implemented the same base-by-base agreement-based accuracy calculation in our previous work **[57]**. This entire part of the study has used 70% of the Dataset “A” as the training set to build the model and its performance on the remaining 30% totally unseen test dataset. **Figure 5** depicts the operation of the implemented deep-learning encoder-decoder system. Further to this, the evaluation was also done through 10-fold random independent train-test trial runs while building the model from the scratch every time maintaining the same 70:30 ratio of train:test datasets to evaluate the consistency of the developed method across variable learning and testing datasets.

### Long read sequencing datasets and related assembling data generation

Long-read sequencing datasets from multiple organisms were retrieved from public repositories like NCBI and Phytozome **[69]**. ONT reads were obtained for six species: *Zea mays* (SRR29027948, SRR29027947), *Glycine max* (SRR16611063, SRR34853577), *Hordeum vulgare* (SRR29563912, SRR29563913), *Oryza sativa* (ERR13336040, ERR13336041, SRR16230871), *A. thaliana* (SRR30315126), and Human (SRR34291213) **(Table S3)**.

Prior to assembly, adapter sequences were identified and removed using Porechop, a tool specifically designed for trimming Oxford Nanopore adapters from long-read sequencing data. Following adapter trimming, reads were subjected to quality-based filtering to remove short and low-quality reads that would otherwise contribute noise to the assembly overlap graph. The processed reads were assembled using the Canu **[19]** assembler, standard approach for assembly of long reads. The genome size estimate was provided to Canu to guide overlap sensitivity parameters and coverage thresholds. Assembly completeness was assessed by computing standard contiguity statistics (N50, L50, total assembly length) using QUAST **[70]**. The following metrics were computed for each assembly as described here:

**i) Number of contigs:** The total number of contigs produced by an assembler reflects the degree of fragmentation of the assembled genome.
**ii) Total length:** The total assembly length is the sum of the nucleotide lengths of all assembled contigs and represents the total amount of genomic sequence recovered by the assembler.
**iii) Longest contig length:** The length of the longest individual contig in the assembly reflects the assembler’s capacity to resolve complex, repeat-rich genomic regions into extended contiguous sequences without introducing structural breaks.
**iv) Misassemblies:** Misassemblies are structural errors in which the internal organisation of an assembled contig is inconsistent with the reference genome, representing the most consequential category of assembly error because they introduce large-scale sequence rearrangements that cannot be corrected by downstream base-level polishing.
**v) Indel Rate per 100 kbp:** The indel rate, expressed as the number of insertion and deletion errors per 100 kilobase pairs of aligned assembly sequence, quantifies the base-level accuracy of the assembly with specific reference to the predominant error type produced by third-generation long-read sequencing platforms. A lower indel rate indicates higher base-level fidelity and reflects the effectiveness of the error correction strategies applied to the raw long reads prior to assembly.
**vi) N50:** The N50 is the length-weighted median contig size, formally defined as the length of the shortest contig in the set of contigs that together account for at least 50% of the total assembly length when contigs are sorted in descending order of length. It is the most widely reported single-value summary of assembly contiguity, providing an intuitive measure of the typical contig size encountered in the assembly.
**vii) Genome Fraction (%):** The genome fraction represents the percentage of the reference genome sequence that is covered by at least one aligned contig from the assembly, providing a direct measure of assembly completeness with respect to the known reference.
**viii) NGA50:** The NGA50 is a reference-informed contiguity metric that represents a more rigorous and biologically meaningful extension of the conventional N50 statistic. NGA50 is calculated from the lengths of aligned contig blocks referred to as Longest Alignment Blocks (LABs) result from breaking assembled contigs at every point of structural misassembly detected upon alignment to the reference genome.

## Results and Discussion

### Data collection and formation of datasets

To develop a generalizable deep learning framework capable of performing error correction and sequence extension across biologically diverse genomes, we curated genomic resources from 16 eukaryotic species encompassing monocots, dicots, woody perennials, basal angiosperms, and animals. Plants were preferred here as they provide much higher genomic variability **[58, 71]**. Whole-genome assemblies, gene annotations (GTF), and protein-coding sequences were retrieved from Ensembl Plants and NCBI as mentioned in the methods section. Across the sixteen species, genome sizes varied substantially from 135 Mb (*A. thaliana*) to 5.1 Gb (*H. Vulgare*) capturing broad genomic complexity that contain 14-85% repeat content (14% *A. thaliana* −85% *Z. mays*) **(Table S4)**. GC content analysis revealed a diversity range of 32.1% to 48.7%, reflecting notable variability in nucleotide composition that is essential for assessing model performance across compositional biases. This diversity spanning genome size, evolutionary distance, promoter organization, and GC content ensures that the trained model is not constrained to narrow sequence domains and is capable of operating across biologically heterogeneous genomic contexts.

2kb upstream promoter regions relative to annotated transcription start sites (TSS), along with their associated protein-coding genic sequences were extracted for *O. sativa*, *Z. mays*, *A. thaliana*, and *V. vinifera.* These sequences collectively formed Dataset ‘A’, comprising 135, 858 promoter-gene pairs across the four species. The remaining 12 species were processed same way to generate Dataset ‘B’, consisting of 332, 896 promoter:genic region pairs. Dataset ‘B’ was reserved exclusively for independent cross-species evaluation and remained strictly unseen during model training, enabling rigorous assessment of out-of-distribution generalization beyond the training species **(Table S2)**.

In addition, the complete genomes of the four primary species were segmented into overlapping 64 bp *k*-mers (explained in the section below) to construct Dataset ‘C’, yielding approximately two million unique sequence fragments. This large-scale *k*-mer dataset served as the foundational input for training the diffusion-based sequence model, allowing the network to learn fine-grained nucleotide dependencies and local sequence statistics essential for accurate error correction. With these datasets established the stage was ready to evolve the desired deep learning architecture to capture nucleotide-level dependency patterns to recover and generate missing bases and their valid sequences.

### Landscape of sequencing errors in nanopore reads with respect to sequence composition and genomic context

To systematically characterize the error profile of nanopore sequencing, we analyzed raw ONT reads from the four representative plant genomes *Oryza sativa, Glycine max, Hordeum vulgare,* and *Zea mays* spanning diverse genome architectures **(Table S3**; data sources: Phytozome **[69]** and NCBI). Following preprocessing, reads were aligned to their respective reference genomes using minimap2 **[72]**, enabling quantitative estimation of insertion, deletion, and substitution errors. Across all the datasets, nanopore reads exhibited substantial but structured error variability, with mean error rates ranging from 12.0% in *Hordeum vulgare* to 24.1% in *Zea mays* **(Figure 6a)**. Error rates were broadly distributed between 5-25%, with a pronounced central tendency around 12-20% (Kruskal-Wallis test, *P* < 1×10⁻¹⁰), indicating a consistent baseline error regime across species.

**Figure 6:**
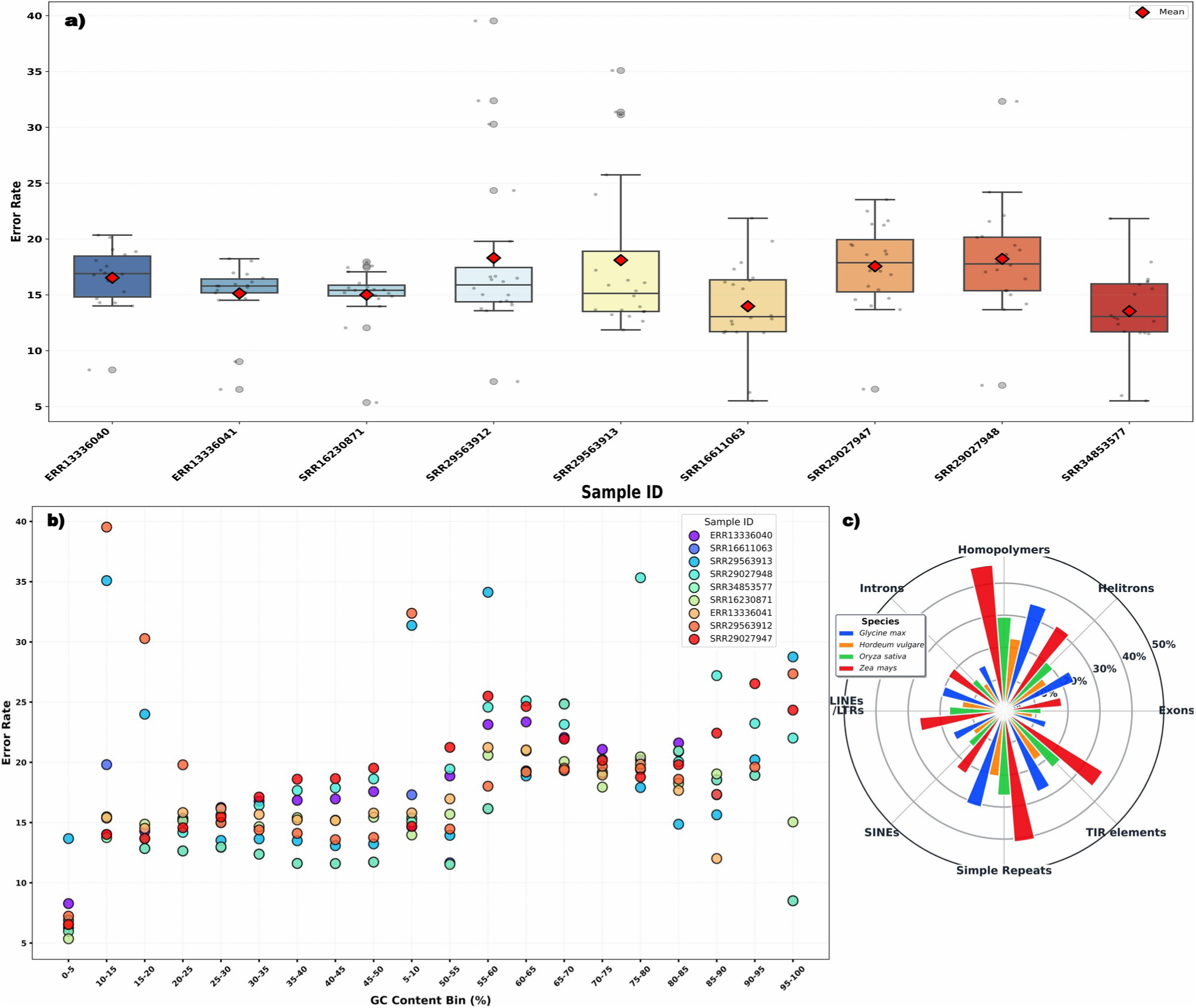
Distribution of ONT sequencing error rates across various genomic sub-elements. **a-b)** Box and scatter plots showing the distribution of error rates (%) across sample files of different species and GC% content bins. **c)** The grouped bar chart illustrates the mean error percentage across four economically significant plant species: *Hordeum vulgare*, *Oryza sativa*, *Glycine max*, and *Zea mays*. Errors are categorized by genomic context, including repetitive regions. Plot reveals a significant error spike in highly repetitive landscapes, particularly in *Zea mays*, whereas unique coding regions maintain higher fidelity across all taxa.

However, this apparent uniformity masked notable sample-specific heterogeneity: while several datasets (e.g., SRR16230871, ERR13336041) displayed tightly constrained error distributions (median ∼15%), others (SRR29563912, SRR29563913) showed marked dispersion, with error rates extending up to ∼40% **(Figure 6a).** These observations suggest that besides specific factors technical and complicated factors also introduce substantial variability in sequencing fidelity.

Study for error distribution at individual reads level reflected that the error rates remained broadly distributed, but a noticeable pattern was observed: 3-23% of reads longer than 10 kb contained high-error-rate sub-sequences (HERS; error rate >50%). Notably, the prevalence of HERS increased with read length (Spearman’s ρ = 0.62, *P* < 1×10⁻⁸) and was significantly higher in ultra-long library preparations compared to standard protocols. These high-error segments represent localized failure modes of nanopore sequencing and have important downstream consequences: error-correction algorithms frequently fragment reads at these regions, thereby eroding the contiguity advantage of long-read sequencing in *de novo* assembly.

### Impact of GC content on error rates

To investigate the determinants of these error patterns, we analyzed the relationship between sequencing accuracy and GC content. A clear U-shaped dependency emerged **(Figure 6b)**, with minimal and stable error rates observed in the balanced GC range (25–55%), where errors consistently remained within 12–18% across all the datasets. In contrast, extreme GC compositions were strongly associated with elevated error rates. AT-rich regions (5–20% GC) exhibited sharp increases in error, reaching up to ∼40%, whereas GC-rich regions (>75%) showed erratic and sample-dependent error spikes, in some cases exceeding 35% (e.g., SRR29027948).

### Regions with the highest incidence of errors

Genomic annotation further resolved error hotspots to specific repetitive elements. Error rates were significantly enriched in simple sequence repeats (SSRs) and DNA transposons compared to unique genomic sequence **(Figure 6c**; P < 10⁻¹⁵, Mann–Whitney test). Homopolymers (≥6 bp) accounted for 41.8% of all indel errors despite comprising only 8.2% of the reference genome. Interestingly, we observed a dichotomy between transposon classes: DNA transposons exhibited higher error rates than retrotransposons (LINEs/SINEs).

Taken together, the analysis revealed that nanopore sequencing errors are not absolutely uniformly distributed but also influenced by an interplay between read length, sequence composition, and genomic context. While a better accuracy regime exists for balanced genomic regions, GC skewed and repetitive regions pose big challenges for nanopore sequencing, leading to direct implications for downstream error correction and genome assembly stages.

### There is a limit to depth based MSA led consensus building

Multiple sequence alignment (MSA) remains a cornerstone of indel error correction in long-read sequencing **[73].** To evaluate the effectiveness of MSA for error correction and consensus building, an analysis of consensus accuracy with respect to increasing read depths was performed. Reads were aligned using MUSCLE **[61] (see Methods section)**, generating MSA blocks in the range of 1X-100X depth. In these MSA blocks positional nucleotide variation could be studied for insertion, deletion, and substitution errors with respect to depth and impact on consensus building.

Across all the datasets, a monotonic improvement was observed for consensus accuracy with increasing coverage **(Figure S1)**. At lower depths (1X-5X), consensus sequences were effectively indistinguishable from individual reads, with indel error rates of ∼15-24% and substitution rates of ∼3–5%. As the coverage increased, consensus accuracy improved significantly with a pronounced transition between 20X and 30X. Majority voting across reads reduced indel rates to ∼8-12% and substitution rates to 1.5-2.8%. This improvement was statistically significant (Spearman’s ρ = −0.81 for indels, −0.76 for substitutions; P < 1×10⁻¹⁰).

Beyond 30X coverage, accuracy gains plateaued, with residual indel rates stabilizing below 10% and substitution rates below 1%. Pairwise comparisons between 30X and higher coverage levels (50×, 100×) showed no significant improvement (Wilcoxon signed-rank test, P > 0.05), establishing ∼30X as a practical threshold for consensus convergence under standard MSA conditions **(Figure S1)**.

Despite this depth-dependent improvement, a persistent and depth-independent failure mode inherent to MSA-based correction was identified. Specifically, a subset of alignment columns exhibited systematic misplaced positionings, in which true nucleotide signals were wrongly placed due to global scoring systems of MSA. Such MSAs are usually not biologically aware and get influenced by indel errors in individual reads **(Figure S1).** This results into incorrect column-wise consensus calling, even when sufficient read depth is available. Quantitatively, 12-18% of consensus positions were affected by such column-shifting artefacts across all coverage levels, with no significant reduction at higher depth (Kruskal–Wallis test, P = 0.27), indicating that this error class is intrinsic to alignment structure. These positions were significantly enriched for indel errors relative to correctly aligned columns (Mann–Whitney test, P < 1×10⁻¹², effect size r = 0.52).

Importantly, these unresolved indel errors had direct consequences for downstream process of genome assembly. Consensus sequences derived from shift-affected regions exhibited localized sequence discontinuities which disrupted overlap detection during assembly, leading to fragmentation of contigs at these positions **(Figure S1)**. This establishes a mechanistic link between MSA column integrity, consensus accuracy, and assembly contiguity, identifying column shifting as the dominant limitation of conventional MSA-based error correction for ONT data.

Another barrier to genome-scale application was computational complexity associated with preparing such MSAs, which are usuaslly progressive. Exhaustive all-versus-all alignment scales results in quadratic time complexity (T) with respect to the number of reads: *T* (*n*)∼*O* (*n*^2^. *L*), where ‘*n’* is the number of reads and ‘*L’* is the average read length. It renders building MSA after best pairs search a highly time consuming step for large datasets. Benchmarking on a moderate dataset (∼500, 000 reads) required >120 hours of time and peaked at 240 GB RAM using 3.2 GhZ clock speed with 256 L3 Mb cache. This computational bottleneck, combined with the inherent accuracy limits of standard MSA, motivated the adoption of an alignment-free pre-clustering strategy. It tremendously accelerated the process of MSA at much lesser resources and time.

### MinHash-based clustering overcomes the time-resource challenge of pairwsie alignment and MSA building

To overcome the inherent limitations of alignment-based read grouping, a MinHash-based locality-sensitive hashing (LSH) strategy was implemented to cluster reads based on sequence similarity prior to MSA. This approach was motivated by the indel-dominated error profile of ONT reads (15–25%), which systematically underestimates true sequence similarity and fragments homologous read groups **[74–76]**. As already mentioned above, to build an MSA one primarily requires to perform all to all similarity search with alignment. A task which becomes impractical for sequencing reads which come in millions. To ensure unbiased clustering across reads of highly variable lengths, ONT reads were fragmented into uniform 1kb segments. This normalization significantly reduced variance in k-mer representation across reads, enabling consistent similarity estimation independent of original read length **[77]**. The 1kb threshold was selected based on standard minimum length requirements in long-read assembly pipelines **[78]**.

Each fragment was represented as a set of overlapping 6-mer shingles, capturing local sequence composition while maintaining robustness to indel perturbations. The interval to see the next indel to occur was around 6 bases **(Figure 2b)**. Thus, 6-mer shingles were considered here to measure the Minhashing similarity between the reads. MinHash signatures spectrum derived from these shingle sets provided a compressed probabilistic representation of sequence content, enabling efficient approximation of Jaccard similarity between reads without explicit alignment **[74].**

Systematic evaluation of five Jaccard similarity thresholds (0.4-0.8) was carried out to identify the optimal balance between cluster purity, consensus accuracy, and read recall **(Table S5)**. Permissive thresholds (0.4–0.5) yielded high recall (>95%) but suffered from low cluster purity (61–75%), resulting in chimeric alignment blocks and elevated consensus indel rates (11.6–13.4%). Conversely, stringent thresholds (0.8) maximized purity (97.1%) but fragmented clusters below the depth required for effective error correction (mean size 14X), paradoxically increasing consensus error to 7.3% due to insufficient redundancy. A threshold of 0.7 emerged as the optimal operating point, achieving 93.6% cluster purity and reducing consensus indel errors to 6.7% at 30X coverage while retaining 93.6% of reads **(Table S5)**.

This approach reduced the computational burden of similarity estimation from *O* ( *n* ² *L*) to approximately *O* ( *n*) under LSH banding schemes, effectively eliminating the need for exhaustive pairwise comparisons and enabling scalable clustering of large ONT datasets.

### Overlap-aware cluster merging restores contiguity without compromising specificity

To further improve cluster structure, a terminal overlap-based merging strategy was implemented, consolidating clusters sharing suffix–prefix overlaps. Systematic evaluation of overlap thresholds (16–256 bp) identified 64 bp as the optimal threshold **(Table S6)**. Short overlaps (≤32 bp) resulted in over-merging of distinct genomic loci, significantly reducing cluster purity (P < 1×10⁻⁵, Mann–Whitney test), whereas longer overlaps (≥128 bp) caused under-merging, fragmenting clusters and reducing effective coverage. At 64 bp, clusters achieved mean sizes of 38 ± 12 reads, consistent with target sequencing depth and enabling robust consensus generation.

### Iterative hexameric walking improves the MSA and subsequent consensus formation

Despite robust clustering, initial multiple sequence alignment (MSA) using MUSCLE **[61]** revealed a persistent structural artifact whose introduction in a previous section was already done: systematic column shifting, where genuine bases were displaced by spurious gaps introduced to accommodate indel errors. This misaligment affected 15.3% of alignment positions and directly disrupted downstream assembly continuity. Analysis of error distribution revealed that ONT indels exhibit statistically significant clustering, with the probability of an error at any position elevated within a ∼6-bases window following a preceding indel. Leveraging this insight, an iterative 6-mer refinement protocol was developed wherein 6-bases blocks were progressively truncated from MSA termini and realigned to resolve edge-case anchoring errors. It was like considering local regions with higher weightage. This countered the impact of globally carried forward indel penalties as MSAs are global alignments where gross similarity diminish the quality of local information. This is the reason that we see errors seeping into the error correction and consensus building steps during assembly.

Quantitative evaluation demonstrated that iterative refinement reduced column-shifted positions from 15.3% to 6.5% at 30X coverage. Overall consensus indel rates decreased from 14.2% to 2.2%, with the most pronounced improvements observed in homopolymeric runs (≥4 bases), where position-specific error rates fell from 19.3% to 2.2%. This reduction in consensus indel rate translated into improved assembler joining rates at cluster boundaries; 91.3% of refined consensus fragments were successfully incorporated into contiguous assembly paths compared to 67.8% of unrefined fragments a 34.7% relative improvement in assembly continuity. This was a huge improvement in assembling just due to this MSA refinemnet step alone.

### Positional noise identification

Despite the substantial improvements achieved through MinHash clustering and iterative MSA refinement, certain positions within the refined alignment blocks inevitably exhibited nucleotide distributions. These positions had no single nucleotide with statistically dominant representation. The fraction of such noisy position within refined MSA blocks across various coverage levels ranged from 38.2% at 5X coverage to 4.7% at 30X coverage, suggesting that the majority of positional uncertainty was depth-resolvable but still a fraction of residual noise persists even at high coverage, representing the most refractory positions in the alignment that neither increased depth nor refined alignment could resolve. Such positions were therefore explicitly designated as noise sites requiring correction by the DDPM.

The corrected MSA blocks in which noise positions were explicitly labelled and spatially registered against their corresponding correct consensus bases thus, constituted the high-fidelity paired training dataset from which the DDPM could now learn the mapping from erroneous alignment tensor representations to biologically accurate consensus sequences.

### Recovering back the corrupted sequences: In the forest of noise, where noise becomes guide

The above mentioned steps ensured a representation of reads in the form of high confidence blocks and errors placed correctly where recovery of correct bases was needed. Such rectified MSA blocks can be seen analogous to images with corrupted pixels equivalent to the errored positions. And this made a fit case of recovery through generative sampling where errors are noise while placing the errored sample at some time-step. From here the lost bases were recoverable while moving back from the given errored time-step. This exact philosophy was applied here through denoising diffusion probabilistic model (DDPM).

A set of such corrected MSA and corresponding reference sequence as target was fed into the DDPM system, during training. A total of 2×10^6^ such MSA blocks from four species (*O. sativa*, *Z. mays*, *A. thaliana*, and *V. vinifera*) to built this dataset. This covered wide range of variability, ranging from 32.1% GC to 48.7% GC. This dataset was divided using a 70:30 training-to-testing split, followed by 10-fold randomized independent train:test trails, ensuring unbiased performance estimates and stability across training cycles. The training set contained 1, 400, 000 MSA blocks while the testing set contained 600, 000 MSA blocks.

### The DDPM architecture

To enable diffusion-based modeling, each nucleotide position in the refined MSA was converted into a structured numerical representation using OHE, while gaps were assigned an all-zero vector to reflect uncertainty. These blocks were encoded into a 64×64×4 three-dimensional tensors using a row-major mapping strategy that preserved local nucleotide adjacency and alignment context within the grid. Within the DDPM framework, these 64×64×4 MSA blocks undergo a forward diffusion process where Gaussian noise is progressively added to the nucleotide states, mimicking the stochastic error processes observed in long-read sequencing while maintaining the original positional alignment of the MSA. By the end of this, it eventually transforms the tensor block into an isotropic noise distribution, providing the training targets for the model. The core component of the DDPM is the Gaussian denoising network, typically implemented as a U-Net. Its was employed to capture hierarchical sequence features across diffusion steps to reconstruct the original aligned block from noise. The encoder processes input vectors to understand the hidden patterns within these blocks, while the decoder uses that understanding to generate corrected sequences step-by-step. This DDPM leverages contextual information from neighboring positions (vertically and horizontally) to implicitly learn conservation patterns and error signatures simultaneously. Once the reverse process generated multiple corrected candidate sequences, they were used to draw a final consensus sequence.

### Initial training and evaluation of the DDPM

The training and evaluation of the DDPM for MSA block correction followed a structured progression, transforming noisy tensors back into coherent meaning sequence through the following steps:

## 1. Training of the DDPM model

During the initial training phases, the model focused on learning the coarse “global patterns” of the DNA sequence layout within the 64×64×4 grid. The loss rate, measured as the difference between proposed and actual noise (Mean Squared Error), showed a steep decline in early epochs as the initial feature width of 128 channels acted as a high-dimensional projection which transformed raw 4-channel DNA nucleotides into 128 channel vector embeddings. These 128 channels allowed the model to capture critical local patterns in the blocks. As the training progressed, the normalization layers stabilized the mean and variance of the outputs, allowing the model to learn the base level patterns for the accurate identification of the bases within the MSA blocks. During the training process, the linear variance scheduler (***β***) dictates the systematic injection of noise into MSA blocks, ensuring distinct noise levels that facilitate precise step-by-step decoding during the reverse diffusion process. A well-tuned ***β*** prevented the model from losing the structural identity of the sequence, ensuring the model was able to distinguish between indels/substitutions and background Gaussian noise.

## 2. Timestep selection and convergence

Initially, the model was configured with 1, 000 timesteps (***T***=1000) to ensure a granular and smooth transition from Gaussian noise to corrected MSA blocks. During the forward process, these 1, 000 steps incrementally corrupted the input tensor until the row-major mapping of nucleotides was completely obscured by noise. In the reverse phase, observations of the model’s convergence revealed that the most significant recovery occured within the first hundred steps. With timestep of 100 **(Figure 7a)** the U-Net aided by its 128-channel initial feature width successfully reconstructed the primary “pixels” or consistent blocks where very few error existed. This stage was sufficient for convergence because the model had already leveraged the context of neighboring positions (vertically and horizontally) to resolve the majority of indels and inconsistent base allocations. Finally, the model produced a corrected representative sequence, allowing for a highly accurate “base-by-base” agreement of average of ∼92% when matched against original target sequences. The accuracy of the corrected sequences ranged from a minimum of 88.86% to a maximum of 93.98%, based on a dataset of 600, 000 MSA blocks totaling ∼32 Mb of sequence information.

**Figure 7:**
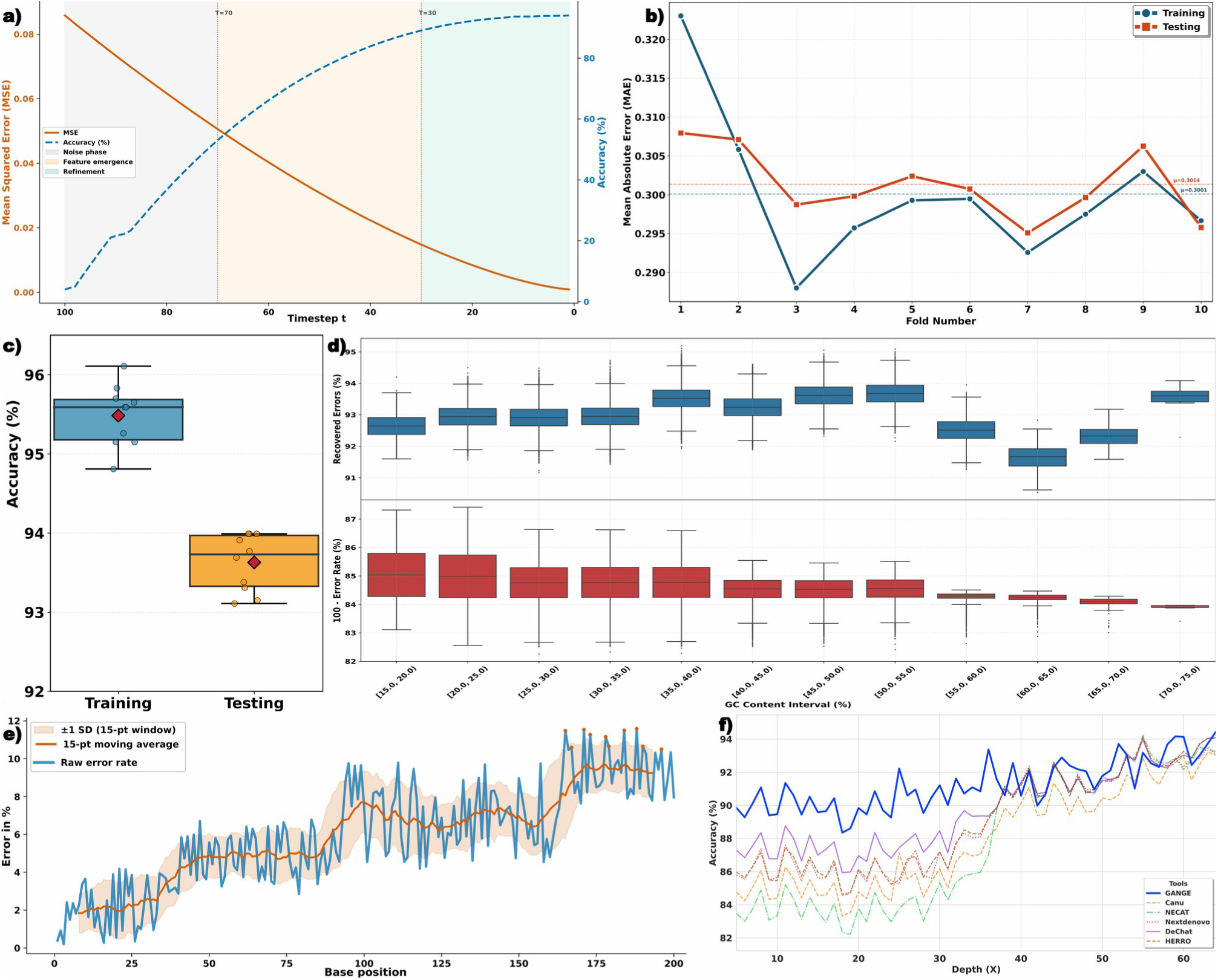
Performance evaluation. **a)** This plot depicts the accuracy of the DDPM model in error correction across different timesteps implemented. **b)** This plot illustrates the training and testing MAE for the DDPM model across 10 random independent train:test trails. The convergence of these values suggests the model’s error rate remains consistent, with MAE values generally fluctuating between 0.285 and 0.325 suggesting highly stable model. **c)** This plot displays the training and testing accuracy for each such trial of DDPM. The DDPM model demonstrates high performance, maintaining a testing accuracy between approximately 93.0% and 94.0%, while training accuracy peaks around 96.0%. **d)** The box plot shows accuracy of DDPM to recover bases that were lost due to poor sequencing quality in ONT read with respect to to GC%. **e)** The line plot shows the accuracy of transformers to generate bases upstream to the given input sequence. **f)** Performance comparison for the varying sequence depth.

## 3. Final parameters of the initial DDPM model

The configuration of the initial DDPM model was optimized to balance the capture of local and global patterns within the 64×64×4 input shape. By projecting MSA blocks into an initial feature width of 128 channels with a channel multiplier of [2, 1, 2, 4] immediately doubled the capacity at the first level to capture high-density local patterns. It finally reached a maximum of 512 channels (128 X 4) at the deepest bottleneck. The core processing power of the network was driven by 6 residual blocks and 4 attention blocks, which work in tandem to preserve the original alignment context while analyzing how all words relate to each other. The residual blocks utilize skip connections to prevent the vanishing gradient problem, ensuring that the “image-like” MSA block remains coherent during deep feature extraction. Meanwhile, the four multi-head attention modules allowed the model to selectively focus on the different sequences within the block. To ensure stable convergence, the model employed a ***β*** schedule of 0.001 and a learning rate of 0.001 using the Swish optimizer. This fixed variance schedule provides a controlled forward diffusion process, preventing the MSA blocks from being corrupted too rapidly and allowing the model to learn specific noise signatures without losing the underlying hidden patterns. Furthermore, weight decay of 0.001 and integrated dropout layers served as critical regularization techniques to combat overfitting, ensuring that the model generalized conservation and contextual pattern rather than simply memorizing training blocks. This initial trained DDPM model with prefixed hyper-parameters achieved an accuracy of ∼92%.

### Hyper-parameter optimization of the DDPM Model led to further improvement

The hyperparameter optimization of the DDPM was structured to balance representational depth with training stability, starting with the configuration of the initial feature width at 128 channels. In early training observations, this width was crucial for capturing critical local patterns and base level information within the 64×64×4 MSA blocks. By utilizing a channel multiplier of [1, 2, 2, 4] after trying several combinations, the network implemented a hierarchical scaling mechanism that doubled processing capacity as spatial resolution decreased. This allowed the model to first analyze local nucleotide adjacency at the 128-channel level and gradually “zoom out” to capture global genomic patterns using up to 512 channels at the deepest layers, effectively preventing over-parameterization while expanding the capacity. To ensure efficient feature refinement and stable gradient propagation, the architecture incorporated two residual blocks per resolution level after exploring multiple combinations of residual blocks. The performance improved significantly when eight self-attention layers were added at intermediate spatial resolutions (16×16 and 8×8), which allowed the model to integrate long-range dependencies and preserve global contextual relationships vital for genomic coherence.

Group normalization with 32 groups was applied throughout the network, providing stable optimization under small-batch training conditions and ensuring consistent feature scaling across diffusion timesteps. The forward diffusion process followed a linear noise variance schedule, with β values ranging from 1×10^-5^ to 0.02 with 1×10^-4^ performing as the best. This schedule enabled gradual corruption of input sequences, allowing the model to learn fine-grained nucleotide corrections at early timesteps and more global sequence restoration at later stages. Gradient clipping (−1.0 to 1.0) was applied to prevent instability during back-propagation.

Model training optimized a mean squared error (MSE) loss between predicted and true noise components, consistent with standard DDPM formulations. Optimization was performed using the Swish optimizer with a learning rate of 1×10^-^⁴ and a weight decay of 1×10^-^⁵, providing effective regularization and stable convergence. The batch size of eight and a diffusion depth of 1, 000 timesteps emerged as the best one. Dynamic learning rate adjustment using ReduceLROnPlateau and early stopping (patience = 10 epochs) were applied to prevent overfitting and frugal computation. These constraints ensured that the model stopped training once the validation loss plateaued, resulting in a finalized DDPM capable of highly accurate sequence reconstruction. The synergy between the 128-channel initial width and the tiered multipliers ensured that the final model resolved indels without losing the underlying “grammar” of the DNA sequence present in the MSA blocks.

The most crucial hyperparameter was found to be the initial feature width (128 channels) combined with the [1, 2, 2, 4] channel multiplier as it defined the model’s “resolution”. The 128-channel projection provided the high-dimensional space necessary to identify the missing base using the contextual information present in the MSA blocks that define syntax of the DNA. As the spatial resolution decreased, the hierarchical scaling doubled the processing capacity, reaching 512 channels at the deepest layers to capture global conservation patterns. This balance ensured the network first mastered local nucleotide adjacency before “zooming out” to resolve indels. Without this specific tiered expansion, the model overfits on local noise and fail to derive the long-range contextual patterns required for a coherent result. Extensive empirical evaluation of architectural variants demonstrated that increasing channel depth or residual block count beyond the selected configuration yielded marginal gains at a substantial computational cost. Similarly, deviation from the selected noise schedule resulted in either oversmoothing or unstable reconstructions. The final configuration therefore represents an empirically validated trade-off between expressiveness and efficiency **(Table S7)**. The optimized model achieved a base-level reconstruction accuracy of 93.91% **(Figure 7b)**, representing an improvement of ∼2% following systematic hyperparameter optimization of the DDPM model.

To further assess performance consistency and generalization capability, the model was evaluated using ten-folds random train-test independent trails. For each run, the dataset was reshuffled and re-partitioned to eliminate partition bias and prevent sequence overlap between training and testing sets. Every time it learned completely afresh from new training set and testing on a new test set. Across all the folds, the mean absolute error (MAE) remained highly stable, with mean training and testing MAE values of 0.3001 and 0.3013, respectively **(Figure 7c; Table S8)**. A paired t-test comparing training and testing MAE across the folds yielded a non-significant difference (p ≈ 0.28), indicating the absence of overfitting and confirming strong generalization capability of the trained model. To avoid any bias towards accuracy reported till now, we had derived a dataset consisting MSA blocks which never was the part of training/testing datasets. This dataset consisted of species like as *H. sapeins*, *B. napus, C. clementina, C. sativus, H. vulgare,* and *S. lycopersicum*.

As was noted previously error distribution had appeared related to extreme GC content. **Figure 7d** provides a visual representation of impact of genomic GC% on ONT read error rates and how effectively DDPM performed across its range. The DDPM based error reduction and sequence recovery worked with equal performance for all GC% range, highlighting its universal learning which can work in any sequence composition context. The percentage of recovered errors by DDPM remained relatively stable in low-to-moderate GC regions, with performance peaking in the 35% to 55% range where median recovery values reach approximately 93.5%. Slight dip in recovery was noted for the range of content (median recovery of 91.7%) which rebound back in the extreme range of 70% to 75%.

Collectively, these results demonstrated that the optimized DDPM exhibited stable and accurate generalization across diverse species, establishing it as a highly reliable tool for error correction.

### Horizontal sequence generation through Transformers

By now it is clear that how sequencing errors and noises can be mitigated by generating bases at the errored positions. We called it vertical sequence generation. But for what these errors happened? To achieve longer sequences which make it easy to perform assembling of complex eukaryotic genomes which are also large ones. The next question was that was it possible to generate sequences upstream of what we had got in the hand and reduce even the horizontal sequence coverage need. This way we were building upon our recent success with generating sequences based on existing sequence information **[57–58]**. Transformer learns long range contexts and associations, which makes them pretty capable to generate bases nearby horizontally expanding the sequences from their terminals.

The test of this capability to profess next ***‘n’*** bases sequence given a sequence, we started with implementation of a Transformer Encoder-decoder system about which we have covered in details in the Methods section. For training, sequence information of 200 bases were considered on whose both ends the target sequences upto 2kb had to be generated. The reasons to choose 200 bases as the anchor sequences had two factors:

**(a)** This is the minimum read length from ONT sequences which get filtered for onward assembling process.
**(b)** This is also the average length of the first exons. This needs to be noted here that one of the purposes for horizontal expansion of the anchored sequences was to decipher the promoter regions given the transcription data of species whose genome is not sequenced.

The sequences were converted into *k*-meric words representations while using 2-mers to 7-mers. Our recent work with similar approach of seeing DNA sequences as language has been highly rewarding **[57–58, 79–80]**. Many recent works have also found the similar success stories with their own language models for DNA sequences **[41–44, 81–88]**. Here, we were attempting a pioneering work of generating next 2kb sequence given a sequence. Each step of generation was based on the contextual grammar of the considered *k*-mers. We were generating next *k*-mer (30mer) for which likelihood support was synergistically being derived from 2-mers to 7-mers encoding grammar from the existing sequence. The attempt to generate continuous 2kb sequences in one go proved highly susceptible to error accumulation. Initial analysis as shown in **Figure 7e** revealed that the generated output maintained a relatively stable and low error rate for the first 30 bases, suggesting a high-fidelity region. However, beyond this threshold, a progressive and continuous rise in the error rate was observed. This decay in accuracy indicated a loss of long-range information with the sequence length. Thus, every generated 30-mer was annexed to the terminal of the foundation sequence, creating new 5’ and 3’ ends. These new ends served as the base for generation of the next 30-mer both the ends, while maintaining the same error rate. Using this approach we had attained 2Kb X 2 sequence generation just using 200 bases of existing sequence, with an accuracy of 81.4%. Hyper-parameter optimization, as described in the next section, improved it further.

Before optimization, the primarily designed Transformer generative decoder fetched a reasonable accuracy of ∼82%. This transformer encoder-decoder composition was a dual-stack system of multi-head self-attention and feed-forward layers. Six encoders were employed which transformed the input *k*-mers into high-dimensional embeddings while using sinusoidal positional encodings to preserve sequence order. Thereafter, six decoders reconstruct the target sequence autoregressively, utilizing masked self-attention to ensure that the generated tokens depend only on previously generated tokens. A specialized encoder-decoder attention layer bridges the two stacks, allowing the decoder to focus on specific contextual features from the input (the anchor sequence). Throughout the network, residual connections and layer normalization stabilize the gradient flow across multiple layers. Finally, a linear layer and softmax function converts the decoder’s output into a probability distribution to determine the most likely next token in the sequence.

Such performance was enough indicative of the promise ahead of generating more accurate upstream/downstream sequences. It was necessary to evaluate how various *k*-mer words were contributing to this generative model. *K*-mer wise ablation analysis revealed that as one moved towards the longer *k*-meric representations. The performance monotonically improved. Sharpest jumps were observed for the transitions from dimeric to pentameric representations. After that, performance saturation started. Also, going beyond 7-mers became computationally exhaustive, which didn’t match the performance gain level. The combination of 5-mers, 6-mers, and 7-mers returned the highest average accuracy of 81.4% while also maintaining computational frugality. The cooperating and information enhancement was evident on using multiple *k*-mer words and representations. **Figure 8a** illustrates the findings for this part in further details.

**Figure 8:**
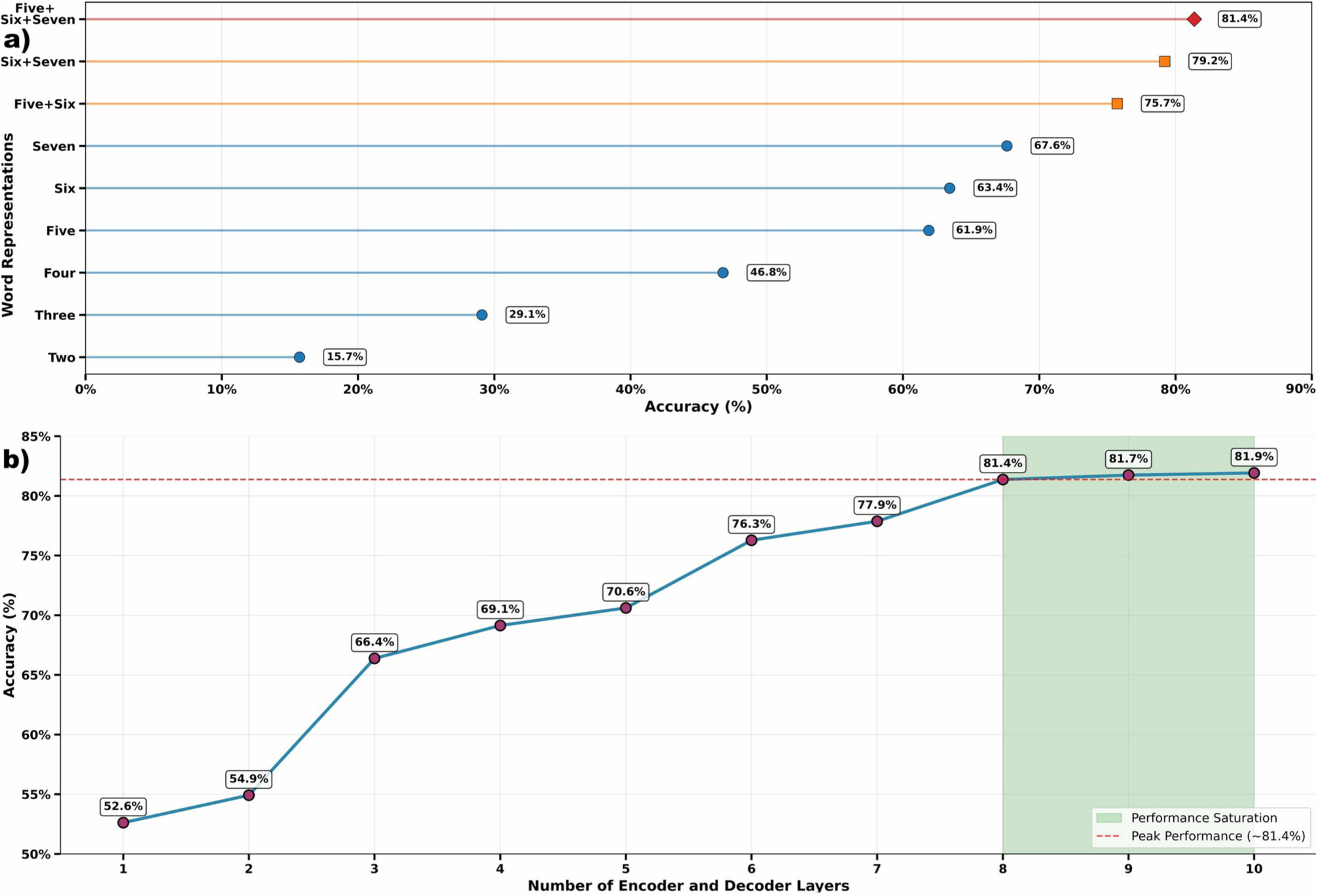
Ablation analysis of k-mer representations, sequence generation, and hyper-parameter optimization of Transformer encoder:decoder pairs. **(a)** Performance of the model across individual and combined k-mer word representations. Accuracy increases progressively with word length, from dimers to heptamers. Combinatorial integration of higher-order k-mers further enhances performance, with five+six+seven achieving the highest accuracy. These results demonstrate that multi-scale k-mer representations captures complementary local and long-range contextual dependencies within the sequences. **(b)** Effect of increasing Transformer encoder–decoder depth on model accuracy. Accuracy rises steadily from 52.6% with one layer to 81.4% at eight layers, after which performance remains consistent, indicating performance saturation. The shaded region highlights the saturation zone beyond eight layers, and the dashed red line marks the peak performance (81.4%). This identifies eight layers as the optimal number for the Transformers.

### Hyper-paramter optimization further improved the transformer model

Eight encoder and decoder layers were found optimal for balancing representational power with a lightweight model architecture **(Figure 8b; Table S9)**. The attention outputs were subjected to a dropout layer (0.12), serving as a regularization mechanism to prevent the model from overfitting and biases. Following the attention mechanism, layer normalization stabilized the input distributions, ensuring consistent gradient flow for deeper layers. The normalized output was then processed by a 42-node feed-forward network (FFN), which acted as a transformation stage to enhance non-linear mapping of genomic features. This enabled the system to extract abstract feature hierarchies like conserved coding regions from the encoded sequences.

A second dropout layer (0.12) and a subsequent 64-node feed-forward layer further expanded the model’s capacity to capture the complex *k*-mer relationship across the sequence variations. In the decoder stage, the model employed a masked multi-head attention mechanism, crucial for the autoregressive generation of flanking sequences. This masking ensured that during the reconstruction of upstream (5′) and downstream (3′) regions, each position only attended to preceding nucleotides, preventing information leakage from future positions. This mechanism is essential for genomic translation tasks where the sequential order of bases determines biological function.

The training strategy utilized cross-entropy loss to penalize deviations between proposed and true nucleotide distributions. Optimization was carried out using the Adam optimizer with an empirically derived learning rate of 0.001 to facilitate stable convergence. Training spanned 100 epochs with a batch size of 6, constrained by GPU memory, and incorporated a weight decay of 1e-4 for L2 regularization. Early stopping with a 10-epoch patience was implemented via PyTorch’s ReduceLROnPlateau, prevented overtraining and ensured the final model achieved an optimal balance between error-correction accuracy, train-test performance gap, and computational efficiency. The final output layer used a softmax activation to produce interpretable probability distributions for the four nucleotide bases, enabling seamless deployment for downstream operations.

The optimized model achieved an accuracy of approximately 84.19%, marking an improvement of ∼2% following hyperparameter optimization. This gain underscored the efficacy of the refined architectural choices, particularly in capturing the non-linear nucleotide patterns essential to generate the unseen immediate sequence while basing on the existing sequence. The training and validation performance curves, illustrated in **Figure 9a-b**, illustrate the convergence behavior and loss/accuracy progression across 100 epochs. The stability of these curves affirmed that the model effectively learned the underlying syntax of the DNA sequences. All the details of hyper-parameters are provided in **Table S10.**

**Figure 9:**
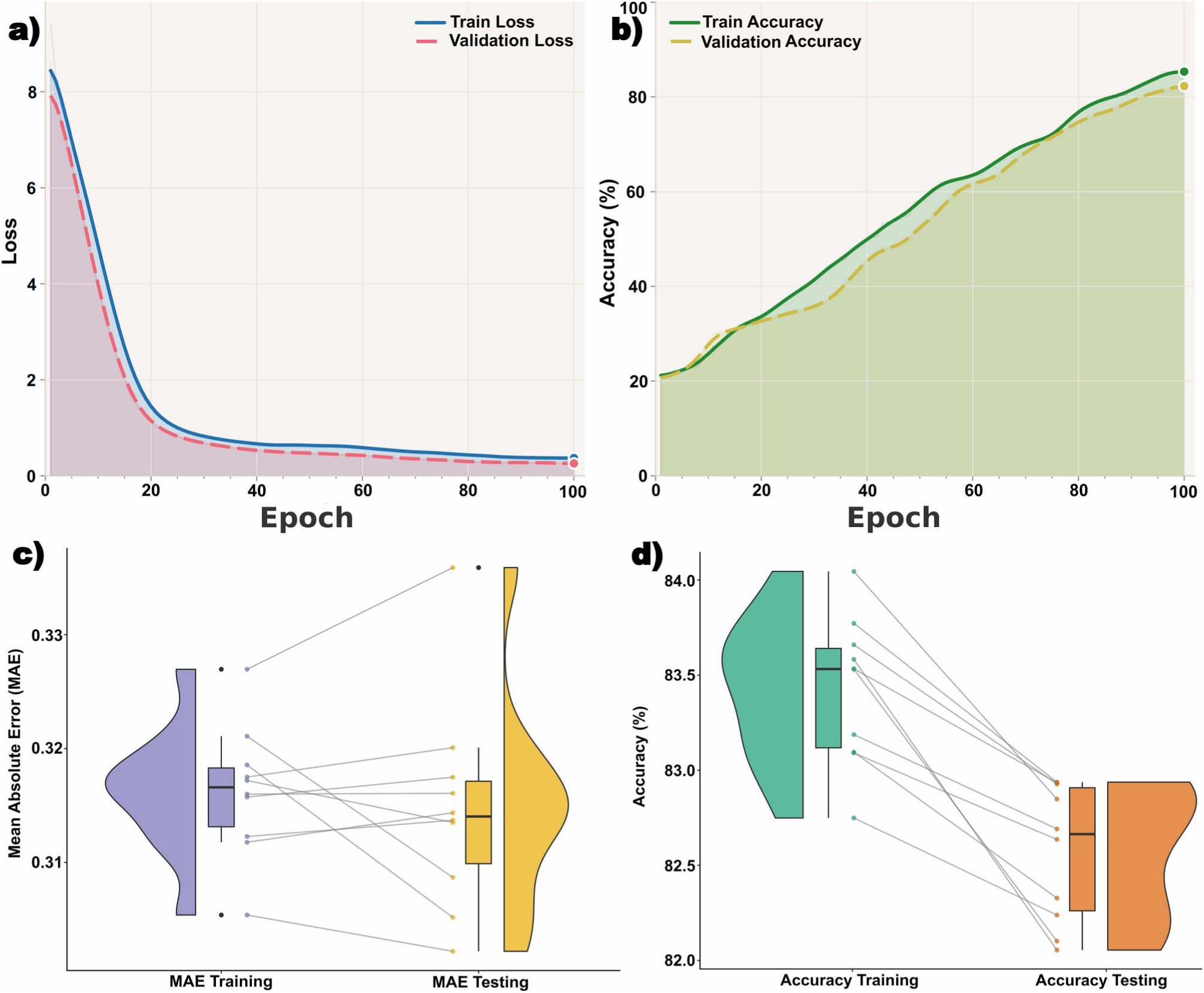
Training dynamics, computational stability, and ten-fold random independent trails performance of the optimized Transformer model. **(a)** Training and validation loss curves across 100 epochs. Both losses decrease rapidly during early epochs and gradually converge, with minimal divergence between training and testing trajectories, indicating stable optimization and absence of overfitting. **(b)** Training and testing accuracy progression over epochs. Accuracy improves steadily for both sets, reaching peak validation accuracy near the final epochs, closely tracking training accuracy. **(c-d)** Ten-fold random independent train:test trails evaluation showing distribution of MAE and accuracy for training and testing sets. Boxplots illustrate minimal variation across folds. The small difference between training and testing MAE and a marginal accuracy gap indicate resistance to overfitting across randomized train:test data splits.

To evaluate accuracy and consistency, the architecture was subjected to 10-fold independent randomized train-test trials. In every iteration, a new set of training and testing datasets was generated with no sequence overlap, ensuring the evaluation remained free from data leakage. The performance trends remained stable, with mean testing MAE values remaining within 0.3150 **(Figure 9d; Table S11)**. The narrow gap between training (MAE) and test MAE indicated that the model maintained a strong generalization capacity. To statistically validate this, a paired t-test between training and test MAE distributions yielded p-values > 0.25, confirming that the model’s performance was consistent and stable. By now, this was clear that through deep-learning a stable horizontal sequence generation system was achievable. It was consistent and stable also. However, there was scope to enhance its accuracy further beyond 84.19%. And for that we returned back to our previously introduced ONT reads error correction system of DDPM.

### Two to Tango: DDPM supported sequence generation by Transformer attains unprecedented accuracy

After hyper-parameter optimization the Transformer encoder:decoder system was generating the 2kb sequences with ∼84% accuracy, the same level of accuracy with which the present generation sequencers like ONT and PacBio generate the reads sequences. And their such accuracy is also highly dependent upon the sequencing depth/coverage. In the previous section on the error correction part, it was clearly demonstrated that how DDPM recovered such high error long-reads from ONT, while improving accuracy level of the corrected reads to ∼94% accuracy, that too at much lower coverage (4X-10X). Thus, if the developed DDPM led vertical sequence generator for error correction was applied on the Transformer generated horizontal sequences, it was guaranteed to attain the final sequences with similar level of accuracy as well as much longer sequences than the actually sequenced sequences. Something very much like sequencing without sequencing. On 200 bases long sequence generated by III^rd^ generation ONT sequencer, we were about to add additional 4kb of sequences with accuracy much more than the sequences generated by the III^rd^ generation sequencers. And very unlike them depending hardly on any high sequencing depth and coverage requirements.

To mitigate the errors in the Transformer generated sequence, the framework employs a secondary, localized DDPM-based correction step. Acting as a “neural polisher”, the DDPM here operates on the OHE representations of the Transformer’s output. Unlike traditional consensus methods that require deep read coverage to identify errors, the DDPM model leverages its learned understanding of biological “DNA grammar” to recognize and reverse non-canonical base transitions and frame-disrupting indels. Every generated 30-mers from Transformer decoder was now a subject input to the DDPM. However, before that these 30-mers would require to be represented as same as the corrected MSA form, which we had introduced previously. Interesting, previously we had MSA from the sequencing reads vertical coverage data, which was not the case here this time. Even such MSA had to be generated now with the transformer generated sequence of 30-mer.

Since, in the back end we would be either having a directly provided corrected sequence or a DDPM corrected consensus sequence to generate the next 30-mers, generation of 30-mers was repeated vertically to mimick sequencing depth. This was done for 100 times, covering the same region with virtual reads generated 100 times for the same region by the transformer part. This mimicked 100X coverage by sequencer which underwent corrective MSA introduced in the previous section. Before introducing the aligned reads in MSA directly to the DDPM module, the MSA was corrected to identify error free and error containing columns, rightfully placed through alignment and corrections. The improved MSA blocks were corrected for missing bases and wrong bases, from which final consensus was drawn to represent the new 30 bases stretch. This newly generated 30 bases was concatenated to the last terminal to generate a new 5’ terminal, on which next 30 bases would be grown. The same was done to the another end of the sequence also, extending the sequencing on both the ends. Through this way the reads were extended 2kb iteratively at both ends, making the fixed sequence much bigger than the actual sequenced read. The generated sequences maintained >92% accuracy, completely on its own without any coverage, and attaining an accuracy higher that what one would achieve through ONT sequencing with high coverage depth **(Figure 10)**.

**Figure 10:**
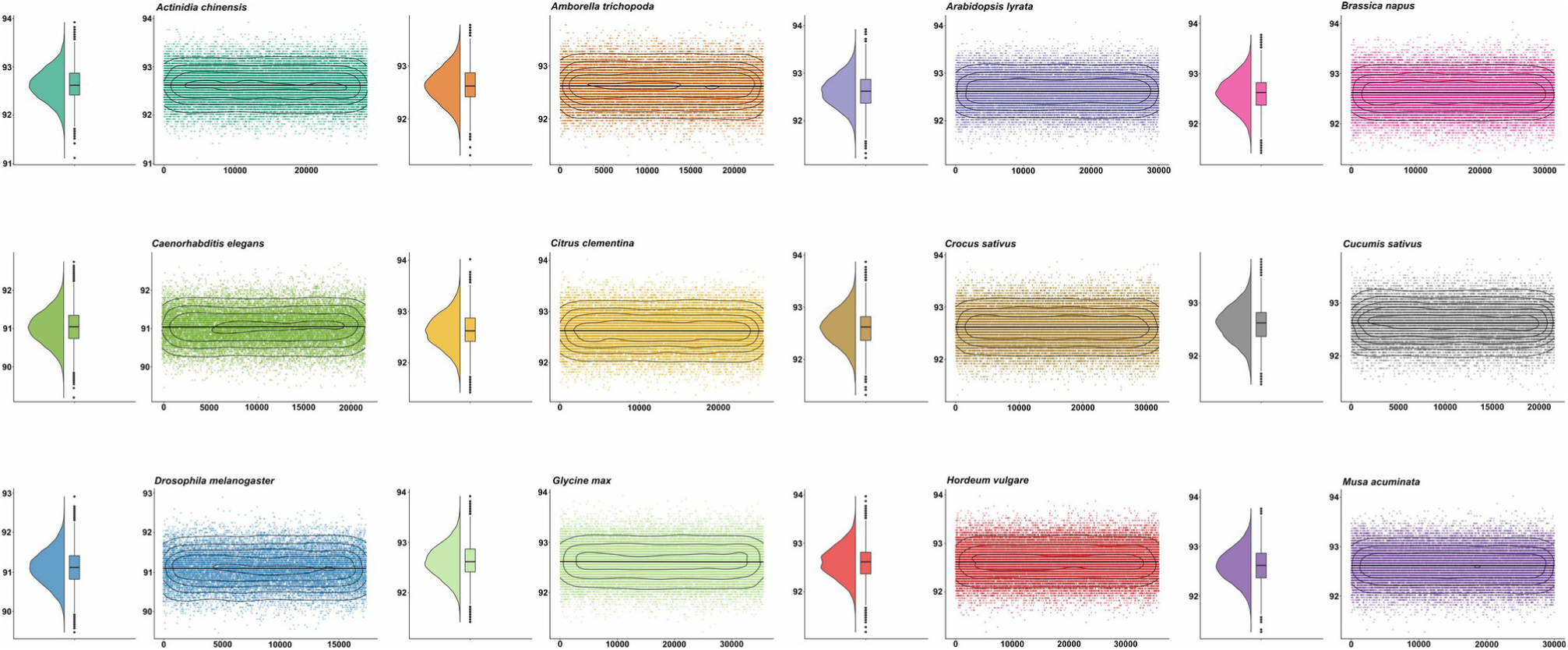
Cross-species performance for promoter generation capability of GANGE. For each species, the left panel displays a violin plot with embedded boxplot summarizing the overall accuracy distribution, while the right panel shows the corresponding scatter-density distribution across sequences. Across all species, accuracy values are tightly clustered around ∼92-93.5%, demonstrating low variance, high reproducibility, and excellent consistency. This summarizes the fact that GANGE can generate highly accurate promoter sequence also if one provides just the 1^st^ exon or transcript data. This makes it possible to perform regulomics research even in those species whose genome is yet not sequenced.

### GANGE can generate upstream promoter regions of unsequenced species

Before moving to the next section two important queries were needed to be satisfied: **(i)** Why the choice of 2kb upstream extension? and **(ii)** can this system be used where the genome is not sequenced?

Both are interconnected queries. Most of species and largely the plant species, are not sequenced for genome. Same time, RNA-seq has become a common practice in such species and the most frequent NGS practice in general. The transcriptome reflects the genic regions. With these genic region sequences alone one can now generate the promoter regions using GANGE. These regions are usually 2kb upstream regions from the gene’s start site. These 2kb upstream sequences host most of the regulatory components like transcription factors binding sites, DNA methylation and epigenetic control switches to regulate the downstream genic region’s expression. Thus, just using GANGE, now one can also carry out regulatory research in species whose genome is not sequenced yet. They can now apply tools like those developed by us on these 2kb regions to decode TFBS, Transcription factor binding regions (TFBRs), condition specific methylome, and condition specific causal nets of regulation **[57–58, 79–80, 89]**.

Thus, we tested GANGE generative system to generate 2kb upstream sequences for 12 species (Dataset B), most of which had no genomic data considered during GANGE training/testing/model making steps. This covered of 332, 896 never seen before genes whose 2kb upstream promoter regions sequences were generated by GANGE. Additionally, we also utilized transcriptome data of saffron and generated 2kb upstream genomic regions of all the transcripts using GANGE. As can be seen in **Figure 10**, highly consistent performance of GANGE was observed for all these species where accuracy remained very close to the one observed over the testing dataset. For all the genes, most of the generated 2kb upstream promoter sequences attained >92% accuracy. This needs to be noted that these species covered a big range of GC content (34%-74%) and represented both monocots and dicots. Also, the observed dispersion of performance was very low for all species.

This study with species whose data were never part of the training and testing steps during the model building of GANGE clearly reinforced its impeccable accuracy in generating sequences. This also underlined the clear possibility to explore the regulomics of all such species whose genome is not sequenced.

### GANGE democratizes low cost genome sequencing by reducing the need of high coverage

An important application-level advantage of the proposed DDPM-based error correction framework is its ability to operate effectively at substantially lower sequencing coverage compared to conventional methods. In standard genome assembly pipelines, accurate error correction typically requires high read depth, commonly >50X coverage, to achieve acceptable consensus accuracy and assembly continuity. At lower coverage (<30X), traditional overlap-layout-consensus or graph-based correctors often show a sharp decline in performance, with residual base-level error rates remaining above 8-12%, leading to fragmented assemblies and misassemblies. To carry out this study, we considered ONT reads from regions different from the genomic regions of *A. thaliana* which were not reflected in the training data of the study. These reads were clustered and binned based on the required depth needed. The compared tools and approaches were ran on this data to get corrected reads which were assembled using Canu **[19]**. Assembled reads were mapped back on the regions using minimap2 **[72]** to evaluate the errors in the sequences.

The GANGE’s approach demonstrated accurate denoising performance at much reduced coverage. The implemented diffusion model can achieve superior correction accuracy even at as low as 10Xcoverage, reducing the required sequencing depth by approximately 55-70% relative to conventional methods. At as low as 4X coverage, GANGE recovered 90% of sequence correctly. To attain the same level, the traditional approaches need at least ∼40X coverage, using the standard and different sequencing and assembly protocols of the compared tools and approaches (**Figure 7f**). At such low coverage levels, the DDPM framework consistently reduced per-base error rates to below 10% **(Figure 7f)**, which is sufficient for smooth downstream assembly and scaffolding. At any given coverage, the sequences recovered by GANGE remained always of better quality than the routine standard approaches. This improvement arises from the model’s ability to learn the underlying sequence distribution and noise characteristics directly, rather than relying on high depth sequencing which is main reasons behind high cost of genome sequencing projects. From application standpoint, this reduction in coverage translates into substantial practical gains. For sequencing large genomes of eukaryotes and plants, which largely remain unsequenced also, one essentially needs >60X coverage at presents. Using GANGE this is possible to achieve the same with 10X coverage, reducing the cost at least by 6-fold, while also drastically reducing the computational resources requirements and time.

### The Aftermath: Superior genome assembly achieved using GANGE

The vertical sequence generation by GANGE reduced the sequencing errors as well as the cost on generating high coverage data. While the horizontal sequence generation of GANGE produced much longer extended reads absolutely without any need of horizontal sequencing and depth. This way two major objectives of any sequencing projects was achieved by GANGE. A natural query arises: How much this all affected the process of genome assembly. To measure this, we introduced GANGE to the standard genome assembling pipeline for long reads using Canu. GANGE was evaluated across assembly of the genomes of *A. thaliana* (SRR30315126), *O. sativa* (ERR13336040 and ERR13336041), and Human chromosome 1 (SRR34291213), all done using ONT long reads alone.

The *Arabidopsis thaliana* sequencing dataset was assembled using GANGE genome assembly workflow (ONT read error correction and extension with GANGE followed by assembly with Canu). Its outcome was compared to the standard default assembly approach (ONT read standard processing and assembly with Canu). The GANGE aided assembling generated a markedly more contiguous assembly, producing only 9 contigs compared to 16 contigs in the default workflow, indicating drastically improved genome assembling with far superior capabilities to resolve complex regions which cause fragmented assembly otherwise. The total assembled genome length increased from 114.2 Mbases to 121.64 Mbases. Additionally, the longest contig got extended to 32.12 Mbases through GANGE, compared to 28.6 Mbases in the default assembly, demonstrating far superior scaffold continuity. Assembly fidelity to the reference genome was also substantially improved.

The GANGE backed assembly emerged more reliable, containing significantly fewer misassemblies (29) compared to the 37 misassemblies found in the traditional assembling approach. This higher accuracy was mirrored in the base-level metrics, where GANGE exhibits a much lower indel rate of 270.98 per 100 kbp, while traditional approach had 472.93 indels per 100 kbp. Interestingly, GANGE actually outperforms the traditional assembling approach with an NA90 of 713, 949 bp compared to 573, 901 bp for the traditional assembling approach. This suggests that GANGE produces longer raw fragments and a more accurate and consistently contiguous representation of the genome. Due to GANGE read extension capabilities three contigs were reduced, which were otherwise missed by the traditional approach. The GANGE led assembly recovered 96.1% complete BUSCO genes, whereas the default assembly recovered 91.4% genes. These metrics demonstrated that GANGE preserved essential gene content with much lesser disruptions within conserved gene families. Overall, the improvements in contiguity, completeness, and reference consistency clearly indicated that GANGE delivered a more accurate and better reconstructed genome of *Arabidopsis thaliana*.

Very similarily, GANGE based *O. sativa* genome assembling produced markedly higher contiguity and accuracy compared to the standard pipeline. Specifically, the number of contigs was reduced from 34 in the default assembly to 20, indicating a significant improvement in sequence continuity and reduced fragmentation. The total assembly length increased slightly to 389.7 Mbases, reflecting better coverage of genomic regions that are often lost or collapsed in standard workflows. A notable enhancement was observed in the longest contig length, where GANGE based assembling achieved 43.2 Mbases, compared to only 35.6 Mbases with the standard method. This suggested superior handling of complex regions, allowing for longer uninterrupted reconstructed chromosomal segments.

Additionally, the GANGE backed assembly shows much lower indel rate of 319.16 per 100 kbp, while the traditional approach had 502.03 indels per 100 kbp. Fewer misassemblies (32) produced by GANGE while the routine default assembly produced 57 misassemblies. Here also, GANGE produced seven less contigs due to its read extension capabilities. Importantly, BUSCO completeness on conserved single-copy orthologs increased from 91.7% in the default assembly to 96.8% using GANGE. This confirms that GANGE backed assembly captures a more complete and biologically meaningful representation of genome and gene space. Moreover, genome fraction (%) to the reference genome improved from 97.83% to 98.12%, further supporting the improved accuracy and reliability of GANGE backed assembly. In both species, the standard approach failed to join several contigs due to its handling of ONT reads where high error rate reads needed better coverage of reads to compensate the errors. It is important to note that for both approaches, insufficient sequencing coverage was a significant factor. Specifically, 1.8% of the genome remained unmapped when the reads were aligned back to the reference, which served as a primary driver for the resulting fragmentation in the assemblies.

The evaluation was further extended to Human Chromosome 1, the largest and most gene-dense human chromosome, often serving as a rigorous test space for error correction and assembly tools. GANGE backed assembly produced an assembly that demonstrated exceptional accuracy to the GRCh38 reference. GANGE’s error correction effectively addressed the Oxford Nanopore-specific indels. A notable enhancement was observed in the longest contig length, where GANGE based assembling achieved 229 Mbases, compared to only 194 Mbases with the standard method. Additionally, GANGE showed much lower indel rate of 210.25 per 100 kbp, while the standard approach returned 254.12 indels per 100 kbp, indicating fewer misassemblies and better agreement with the reference by GANGE backed assembly. **Table S12-13** summarizes the observed performance gain with GANGE enabled genomic assemblies when compared with the standard assembling protocol.

### Comparative benchmarking with existing assembly protocols establishes GANGE as the outstanding one

The selection of an assembly pipeline is a critical determinant of contiguous sequence reconstruction and the base resolution of genomes. To evaluate the efficacy of GANGE, we benchmarked four state-of-the-art (SOTA) read correction strategies for long-read assembly i.e., HERRO **[54]**, NECAT **[90]**, deChat (*De novo* Consensus-based Haplotype Assembler) **[91]**, and NextDenovo **[92]**, focusing on their algorithmic performance and accuracy on *A. thaliana* genome assembly using SRR30315126 read data. HERRO represents a paradigm shift toward DL-integrated assembly. By employing a transformer-based architecture, HERRO performs high-fidelity error correction on noisy ONT reads, effectively “rescuing” sequences that traditional statistical filters would categorize as unalignable. The second software tool, NECAT utilizes a highly optimized “correct-then-assemble” framework based on the Overlap-Layout-Consensus (OLC) strategy. By integrating bit-vector acceleration for rapid overlap detection, NECAT achieves a superior balance between contiguity. For diploid organisms characterized by high heterozygosity, deChat enables repeat- and haplotype-aware error correction, leveraging the strengths of both *de* Bruijn graphs and variant-aware multiple sequence alignment to create a synergistic approach. The last tool included was NextDenovo **[92]** which utilizes a string-graph framework and a specialized 128-bit seed system, which claims to be fastest assembler and with the lowest memory footprint.

To evaluate the performance of software tools benchmarked, we used QUAST **[70]** to derive all the statistics with respect to to assembled genome assembly. The benchmarking results (**Table S14**) clearly identified GANGE backed assembly as the top-performing one, outperforming the compared tools assembly and strategies across all critical QUAST parameters. The most significant advantage was observed in the Genome Fraction, where GANGE backed assembly successfully recovered 98.3% of the reference genome. This indicated a superior ability to navigate the complex, repetitive regions characteristic of completely bug genomes, where other tools like NECAT and DeChat often fail to resolve overlaps, leading to excluded sequences.

GANGE’s assembly delivered N50 value of 23.2 Mb. By achieving a higher N50 with fewer contigs, GANGE demonstrated superior long-range contiguity. From an accuracy perspective, GANGE maintained a substantial lead in base-level precision. With only 270.98 indels counts per 100kb, it significantly reduces the post-assembly polishing burden compared to NECAT and NextDenovo. The comparative evaluation of five genome assembling strategies with GANGE, HERRO, NECAT, DeChat, and NextDenovo revealed substantial differences in assembly quality, accuracy, and completeness on *A. thaliana* (SRR30315126). Among all assemblers tested, GANGE backed genome assembly consistently demonstrated far superior performance across all the metrics (**Table S14**), establishing itself as the most reliable approach for generating high-quality genome assemblies.

In contrast, NECAT and DeChat backed assembly exhibited comparatively lower performance across all the key metrics. NECAT backed assembly generated the highest number of contigs (14) and the lowest total assembly length (118.65 Mbp), suggesting challenges in capturing full genomic content. More notably, NECAT backed assembly had the second highest indel rate (305.93 per 100 kbp), reflecting reduced consensus accuracy. DeChat backed assembly, while producing a moderate number of contigs (12) and total length (120.38 Mbp), also displayed a relatively low indel rate (297.23 per 100 kbp). These findings indicate that while both assemblers are capable of generating contiguous assemblies, they may require additional polishing steps to achieve accuracy levels suitable for reference-quality genome reconstruction.

HERRO and NextDenovo backed assembly occupied an intermediate position in the performance hierarchy. HERRO backed assembly ranked second in accuracy with an indel rate of 287 per 100 kbp and achieved the second-highest genome fraction (97.2%), demonstrating its effectiveness in balancing contiguity and correctness. However, its contiguity metrics, including N50 (22.85 Mbp) and total contig count (12), were slightly inferior to those of GANGE backed assembly. NextDenovo backed assembly produced the longest individual contig (31.90 Mbp) and performed comparably in terms of N50 (22.67 Mbp), yet its indel rate was substantially higher (306.24 per 100 kbp), suggesting that while it excels in generating long contigs, it does so at the cost of sequence accuracy.

The variation in genome fraction which represents the percentage of the reference genome sequence that is covered by at least one aligned contig from the assembly (96.1% to 98.3%) and indel rates indicates differences in how successfully each assembler reconstructed the complex regions. GANGE’s superior genome fraction implies better handling of such challenging regions, likely through Transformer-DDPM based read extension algorithm. In case of all running tools, one point should be noted that the lesser sequencing read coverage to genome played a crucial part. ∼2% part was missing especially in Chromosome 1, 2, and 3 when reads were mapped back to the reference genome. In such scenario too GANGE was able to extend the assembly to join the contigs due to its read extension prowess upto 5kb of completely unsequenced regions other approaches completely failed to do anything here.

These results focus on GANGE prowess which employs effective DDPM based error correction with read extension, enabling it to resolve repetitive and complicated regions more accurately while minimizing assembly fragmentation and same time delivering much more complete assembly at far lesser cost. Overall, this comparative study demonstrated that GANGE backed assembly consistently outperforms the SOTA assembly strategies in terms of contiguity, accuracy, and completeness. This all makes it an indispensible choice for ONT/III^rd^ Gen sequencers based genome sequencing projects where drastic cost cutting is now assured with its use.

### Conclusion

The present work is first ever of its type where clear demonstration has been made about how one can achieve sequencing without sequencing of genome with bare minimum input of low coverage sequencing data. That too from the cheapest and most portable sequencing platforms like ONT, which are otherwise full of indel errors. By applying GANGE on low coverage indel prone reads, one can now build for superior genome. Contigs and scaffolds at as low as 4X to 10X coverage. To achieve the same level, the existing technologies have to go for manifold higher coverage. And despite increasing the coverage, the existing protocols can only match the error correction capabilities of GANGE at much higher coverage. They are absolutely nowhere when extension and mergence of reads for longer coherent assemblies in the form of contigs and scaffolds are used. For that, where the existing protocols essentially require horizontal coverage of genome sequencing and techniques like additional long-read sequencing from multiple platforms/replicates and mate-pair sequencing etc, GANGE gives absolute freedom from all these. GANGE extends both ends of any given sequence by 2kb (4kb in total) without needing any additional experiments, and absolutely based upon the existing sequence. It needs just 200 bases of such anchor sequence and adds additional 4kb to it with >92% accuracy enough to raise big contigs at much lower cost.

This way GANGE performs something like sequencing without sequencing. At much lower coverage it recovers the actual sequences from far cheaper but error prone long reads, replacing the high depth vertical sequencing. And same time it horizontally expands them, mimicking the horizontal genome sequencing.

GANGE achieves this all by first time introduced Diffusion based deep-learning which recovers the correct bases at noisy errored positions, drastically reducing the need of high depth sequencing. And using self attention over genomic language syntax while learning from wide range of genomics data it has raised a robust transformer language model. The Transformer language model is generative one which iteratively speaks out the next 30-mers based on the existing sequence and context hidden in it. The error in the generated 30-mers are in term rectified by the Diffusion modeling part.

Thus, highly accurate sequence is restored from it. This way, this dual system generates the 2kb sequence at both terminals and achieves sequencing without sequencing, both vertical and horizontal way.

Use of GANGE is not just for making genome sequencing now possible at much lesser cost and time. The study has also demonstrated that how accurately it recovered the 2kb upstream promoter regions when just the protein coding regions/1st exons/transcripts were provide. Thus, using GANGE one can now even perform regulomics studies even in those species where genome is not sequenced yet. Otherwise, one essentially requires sequenced genomes to perform gene regulation studies. GANGE now brings freedom at this part also.

GANGE is freely available. And we now expect to see a sudden jump in genome sequencing projects at much lesser costs. Even modest labs with just one handy and small ONT sequencers would now be capable of sequencing an entire eukaryotic genome at far lesser cost. And those who still don’t wish to sequence the genomes, can decode the regulomics of their species of interest with just transcriptomes.

The next objective of GANGE from here is now to make it possible to recover correct sequences from error prone sequencing reads at just 1X coverage while expanding the horizontal coverage from the existing limit of 4kb. And what we have shown here possible, this objective becoming a reality is not distant!

## Acknowledgements

The work was carried out under the aegis of The Himalayan Centre for High-throughput Computational Biology (HiCHiCoB), a BIC supported by DBT, Govt. of India. SG and AK are thankful to DBT, India for financial support as project associateship. UB is thankful for DBT SRF fellowship. SG and UB are also thankful to Academy of Scientific and Innovative Research (AcSIR) for their Ph.D. enrollment. All authors are thankful to the Director, CSIR-IHBT, for his kind support for this study. This MS has CSIR-IHBT MSID ##.

## Author’s contributions

SG carried out the major parts of this study. AK developed the web-server of the GANGE. UB and AK assisted in the study. RS conceptualized, designed, analyzed, and supervised the entire study. SG, UB, and RS wrote the MS.

## Software and Data availability

The GANGE is freely accessible as a webserver at https://scbb.ihbt.res.in/gange/, enabling light-weight task such as promoter generation through an interactive browser-based interface. The platform is implemented using HTML5, CSS3, JavaScript, and PHP. All outputs can be downloaded directly from the web-server. For large-scale analyses, including genome assembly and high-throughput processing, the software is available as a standalone implementation via GitLab (https://gitlab.com/scbblab/gange/). The standalone version supports flexible deployment in local and high-performance computing environments. Comprehensive documentation, installation guidelines, and step-by-step tutorials are provided at the webserver tutorial page, facilitating use and integration into existing workflows. All the secondary data used in the present study were publicly available and their due references and sources have been provided in **Table S1-14**.

## Supplementary Information

**Table S1: List of some published tools which are used at various stages of the sequencing and assembly pipelines.**

**Table S2: Dataset description of collected promoter-gene pairs.**

**Table S3: Details of ONT reads collected from SRA.**

**Table S4: Repeat content of species.**

**Table S5: Evaluation of MinHash clustering performance across Jaccard similarity thresholds using 6-mer shingling on ONT read data.**

**Table S6: Evaluation of terminal overlap length on cluster merging performance and downstream consensus accuracy at a fixed Jaccard similarity threshold of 0.7.**

**Table S7: List of hyperparameter considered for the optimization of the DDPM model.**

**Table S8: MAE of ten-fold random independent train:test trails of the DDPM model.**

**Table S9: Accuracy across different number of encoder and decoder layers of the Transformer model.**

**Table S10: List of hyperparameter considered for the optimization of the Transformer model.**

**Table S11: MAE of ten-fold random independent train:test trails of the Transformer model.**

**Table S12: Statistics of the assembly generated with Standard genome protocol and GANGE backed genome assembly.**

**Table S13: Statistics of the assembly generated with Standard genome protocol and GANGE backed genome assembly.**

**Table S14: Statistics of the assembly generated with different approach backed genome assembly.**

**Figure S1: Error rate on different depths of sequencing coverage before and after MSA refinement.** Plot depicting the relationship between sequencing coverage depth (X) and error rate (%). Error rates decline sharply with increasing coverage from 1X to 50X, followed by a gradual stabilization beyond 50X coverage. This inverse relationship demonstrates that higher sequencing depth improves consensus accuracy by enabling better discrimination between true biological variants and sequencing artifacts.

**Figure S2: GPU utilization during training and validation phases of the Transformer system.** Training maintains consistently high GPU utilization (∼95-99%), while testing requires substantially lower requirement (∼10–20%), confirming frugal application overhead.

## References

1. Dida, F. and Yi, G. (2021) Empirical evaluation of methods for de novo genome assembly. PeerJ Comput Sci, 7, e636.

2. Koren, S. and Phillippy, A.M. (2015) One chromosome, one contig: complete microbial genomes from long-read sequencing and assembly. Curr Opin Microbiol, 23, 110–120.

3. Amarasinghe, S.L., Su, S., Dong, X., Zappia, L., Ritchie, M.E. and Gouil, Q. (2020) Opportunities and challenges in long-read sequencing data analysis. Genome Biol, 21, 30.

4. Stoler, N. and Nekrutenko, A. (2021) Sequencing error profiles of Illumina sequencing instruments. NAR Genom Bioinform, 3, lqab019.

5. Rhoads, A. and Au, K.F. (2015) PacBio Sequencing and Its Applications. Genomics Proteomics Bioinformatics, 13, 278–289.

6. Jain, M., Olsen, H.E., Paten, B. and Akeson, M. (2016) The Oxford Nanopore MinION: delivery of nanopore sequencing to the genomics community. Genome Biol, 17, 239.

7. Aury, J.-M., Engelen, S., Istace, B., Monat, C., Lasserre-Zuber, P., Belser, C., Cruaud, C., Rimbert, H., Leroy, P., Arribat, S., et al. (2022) Long-read and chromosome-scale assembly of the hexaploid wheat genome achieves high resolution for research and breeding. Gigascience, 11, giac034.

8. Faulk, C. (2023) De novo sequencing, diploid assembly, and annotation of the black carpenter ant, Camponotus pennsylvanicus, and its symbionts by one person for $1000, using nanopore sequencing. Nucleic Acids Res, 51, 17–28.

9. Kress, W.J., Soltis, D.E., Kersey, P.J., Wegrzyn, J.L., Leebens-Mack, J.H., Gostel, M.R., Liu, X. and Soltis, P.S. (2022) Green plant genomes: What we know in an era of rapidly expanding opportunities. Proc Natl Acad Sci U S A, 119, e2115640118.

10. Pucker, B., Irisarri, I., de Vries, J. and Xu, B. (2022) Plant genome sequence assembly in the era of long reads: Progress, challenges and future directions. Quant Plant Biol, 3, e5.

11. Huddleston, J., Ranade, S., Malig, M., Antonacci, F., Chaisson, M., Hon, L., Sudmant, P.H., Graves, T.A., Alkan, C., Dennis, M.Y., et al. (2014) Reconstructing complex regions of genomes using long-read sequencing technology. Genome Res, 24, 688–696.

12. Michael, T.P. and VanBuren, R. (2020) Building near-complete plant genomes. Curr Opin Plant Biol, 54, 26–33.

13. Loman, N.J., Quick, J. and Simpson, J.T. (2015) A complete bacterial genome assembled de novo using only nanopore sequencing data. Nat Methods, 12, 733–735.

14. Chin, C.-S., Peluso, P., Sedlazeck, F.J., Nattestad, M., Concepcion, G.T., Clum, A., Dunn, C., O’Malley, R., Figueroa-Balderas, R., Morales-Cruz, A., et al. (2016) Phased diploid genome assembly with single-molecule real-time sequencing. Nat Methods, 13, 1050–1054.

15. Jain, M., Koren, S., Miga, K.H., Quick, J., Rand, A.C., Sasani, T.A., Tyson, J.R., Beggs, A.D., Dilthey, A.T., Fiddes, I.T., et al. (2018) Nanopore sequencing and assembly of a human genome with ultra-long reads. Nat Biotechnol, 36, 338–345.

16. Carneiro, M.O., Russ, C., Ross, M.G., Gabriel, S.B., Nusbaum, C. and DePristo, M.A. (2012) Pacific biosciences sequencing technology for genotyping and variation discovery in human data. BMC Genomics, 13, 375.

17. Jain, M., Fiddes, I.T., Miga, K.H., Olsen, H.E., Paten, B. and Akeson, M. (2015) Improved data analysis for the MinION nanopore sequencer. Nat Methods, 12, 351–356.

18. Ashton, P.M., Nair, S., Dallman, T., Rubino, S., Rabsch, W., Mwaigwisya, S., Wain, J. and O’Grady, J. (2015) MinION nanopore sequencing identifies the position and structure of a bacterial antibiotic resistance island. Nat Biotechnol, 33, 296–300.

19. Koren, S., Walenz, B.P., Berlin, K., Miller, J.R., Bergman, N.H. and Phillippy, A.M. (2017) Canu: scalable and accurate long-read assembly via adaptive k-mer weighting and repeat separation. Genome Res, 27, 722–736.

20. Kolmogorov, M., Yuan, J., Lin, Y. and Pevzner, P.A. (2019) Assembly of long, error-prone reads using repeat graphs. Nat Biotechnol, 37, 540–546.

21. Chin, C.-S., Alexander, D.H., Marks, P., Klammer, A.A., Drake, J., Heiner, C., Clum, A., Copeland, A., Huddleston, J., Eichler, E.E., et al. (2013) Nonhybrid, finished microbial genome assemblies from long-read SMRT sequencing data. Nat Methods, 10, 563–569.

22. Li, H. (2013) Aligning sequence reads, clone sequences and assembly contigs with BWA-MEM. 10.48550/arXiv.1303.3997.

23. Langmead, B. and Salzberg, S.L. (2012) Fast gapped-read alignment with Bowtie 2. Nat Methods, 9, 357–359.

24. Walker, B.J., Abeel, T., Shea, T., Priest, M., Abouelliel, A., Sakthikumar, S., Cuomo, C.A., Zeng, Q., Wortman, J., Young, S.K., et al. (2014) Pilon: an integrated tool for comprehensive microbial variant detection and genome assembly improvement. PLoS One, 9, e112963.

25. Zimin, A.V. and Salzberg, S.L. (2020) The genome polishing tool POLCA makes fast and accurate corrections in genome assemblies. PLoS Comput Biol, 16, e1007981.

26. Morisse, P., Marchet, C., Limasset, A., Lecroq, T. and Lefebvre, A. (2021) Scalable long read self-correction and assembly polishing with multiple sequence alignment. Sci Rep, 11, 761.

27. Luo, X., Kang, X. and Schönhuth, A. (2022) VeChat: correcting errors in long reads using variation graphs. Nat Commun, 13, 6657.

28. Au, K.F., Underwood, J.G., Lee, L. and Wong, W.H. (2012) Improving PacBio long read accuracy by short read alignment. PLoS One, 7, e46679.

29. Hackl, T., Hedrich, R., Schultz, J. and Förster, F. (2014) proovread: large-scale high-accuracy PacBio correction through iterative short read consensus. Bioinformatics, 30, 3004–3011.

30. Goodwin, S., Gurtowski, J., Ethe-Sayers, S., Deshpande, P., Schatz, M.C. and McCombie, W.R. (2015) Oxford Nanopore sequencing, hybrid error correction, and de novo assembly of a eukaryotic genome. Genome Res, 25, 1750–1756.

31. Haghshenas, E., Hach, F., Sahinalp, S.C. and Chauve, C. (2016) CoLoRMap: Correcting Long Reads by Mapping short reads. Bioinformatics, 32, i545–i551.

32. Lee, H., Gurtowski, J., Yoo, S., Marcus, S., McCombie, W.R. and Schatz, M. (2014) Error correction and assembly complexity of single molecule sequencing reads. 10.1101/006395.

33. Bao, E. and Lan, L. (2017) HALC: High throughput algorithm for long read error correction. BMC Bioinformatics, 18, 204.

34. Salmela, L. and Rivals, E. (2014) LoRDEC: accurate and efficient long read error correction. Bioinformatics, 30, 3506–3514.

35. Miclotte, G., Heydari, M., Demeester, P., Rombauts, S., Van de Peer, Y., Audenaert, P. and Fostier, J. (2016) Jabba: hybrid error correction for long sequencing reads. Algorithms Mol Biol, 11, 10.

36. Wang, J.R., Holt, J., McMillan, L. and Jones, C.D. (2018) FMLRC: Hybrid long read error correction using an FM-index. BMC Bioinformatics, 19, 50.

37. Firtina, C., Bar-Joseph, Z., Alkan, C. and Cicek, A.E. (2018) Hercules: a profile HMM-based hybrid error correction algorithm for long reads. Nucleic Acids Res, 46, e125.

38. Eraslan, G., Avsec, Ž., Gagneur, J. and Theis, F.J. (2019) Deep learning: new computational modelling techniques for genomics. Nat Rev Genet, 20, 389–403.

39. Libbrecht, M.W. and Noble, W.S. (2015) Machine learning applications in genetics and genomics. Nat Rev Genet, 16, 321–332.

40. Ji, Y., Zhou, Z., Liu, H. and Davuluri, R.V. (2021) DNABERT: pre-trained Bidirectional Encoder Representations from Transformers model for DNA-language in genome. Bioinformatics, 37, 2112–2120.

41. Ji, Y., Zhou, Z., Liu, H. and Davuluri, R.V. (2021) DNABERT: pre-trained Bidirectional Encoder Representations from Transformers model for DNA-language in genome. Bioinformatics, 37, 2112–2120.

42. Sanabria, M., Hirsch, J., Joubert, P.M. and Poetsch, A.R. (2024) DNA language model GROVER learns sequence context in the human genome. Nat Mach Intell, 6, 911–923.

43. Dalla-Torre, H., Gonzalez, L., Mendoza-Revilla, J., Lopez Carranza, N., Grzywaczewski, A.H., Oteri, F., Dallago, C., Trop, E., de Almeida, B.P., Sirelkhatim, H., et al. (2025) Nucleotide Transformer: building and evaluating robust foundation models for human genomics. Nat Methods, 22, 287–297.

44. Nguyen, E., Poli, M., Faizi, M., Thomas, A., Birch-Sykes, C., Wornow, M., Patel, A., Rabideau, C., Massaroli, S., Bengio, Y., et al. (2023) HyenaDNA: Long-Range Genomic Sequence Modeling at Single Nucleotide Resolution. 10.48550/arXiv.2306.15794.

45. Boža, V., Brejová, B. and Vinař, T. (2017) DeepNano: Deep recurrent neural networks for base calling in MinION nanopore reads. PLoS One, 12, e0178751.

46. Teng, H., Cao, M.D., Hall, M.B., Duarte, T., Wang, S. and Coin, L.J.M. (2018) Chiron: translating nanopore raw signal directly into nucleotide sequence using deep learning. Gigascience, 7, giy037.

47. Boža, V., Perešíni, P., Brejová, B. and Vinař, T. (2020) DeepNano-blitz: a fast base caller for MinION nanopore sequencers. Bioinformatics, 36, 4191–4192.

48. Xu, Z., Mai, Y., Liu, D., He, W., Lin, X., Xu, C., Zhang, L., Meng, X., Mafofo, J., Zaher, W.A., et al. (2021) Fast-bonito: A faster deep learning based basecaller for nanopore sequencing. Artificial Intelligence in the Life Sciences, 1, 100011.

49. Choudhury, O., Chakrabarty, A. and Emrich, S.J. (2018) HECIL: A Hybrid Error Correction Algorithm for Long Reads with Iterative Learning. Sci Rep, 8, 9936.

50. Wang, R. and Chen, J. (2023) RNNHC: A hybrid error correction algorithm for long reads based on Recurrent Neural Network. 10.21203/rs.3.rs-3309460/v1.

51. Wang, R. and Chen, J. (2024) DeepCorr: a novel error correction method for 3GS long reads based on deep learning. PeerJ Comput. Sci., 10, e2160.

52. Wang, R. and Chen, J. (2024) NmTHC: a hybrid error correction method based on a generative neural machine translation model with transfer learning. BMC Genomics, 25, 573.

53. Mastoras, M., Asri, M., Brambrink, L., Hebbar, P., Kolesnikov, A., Cook, D.E., Nattestad, M., Lucas, J., Won, T.S., Chang, P.-C., et al. (2024) Highly accurate assembly polishing with DeepPolisher. bioRxiv, 10.1101/2024.09.17.613505.

54. Stanojević, D., Lin, D., Nurk, S., Sessions, P.F. de and Šikić, M. (2024) Telomere-to-Telomere Phased Genome Assembly Using HERRO-Corrected Simplex Nanopore Reads. 10.1101/2024.05.18.594796.

55. Boetzer, M. and Pirovano, W. (2014) SSPACE-LongRead: scaffolding bacterial draft genomes using long read sequence information. BMC Bioinformatics, 15, 211.

56. English, A.C., Richards, S., Han, Y., Wang, M., Vee, V., Qu, J., Qin, X., Muzny, D.M., Reid, J.G., Worley, K.C., et al. (2012) Mind the gap: upgrading genomes with Pacific Biosciences RS long-read sequencing technology. PLoS One, 7, e47768.

57. Gupta, S., Kumar, A., Kesarwani, V., Bhati, U. and Shankar, R. (2025) DMRU: generative deep learning to unravel condition-specific cytosine methylation in plants. Brief Bioinform, 26, bbaf579.

58. Gupta, S., Jyoti, Bhati, U., Kesarwani, V., Sharma, A. and Shankar, R. (2025) PTF-Vāc: An explainable and generative deep co-learning encoder–decoder system for ab initio discovery of plant transcription factor binding sites. Plant Communications, 6, 101543.

59. Broder, A.Z. (1997) On the resemblance and containment of documents. In Proceedings. Compression and Complexity of SEQUENCES 1997 (Cat. No.97TB100171).pp. 21–29.

60. Jaccard, P. (1902) Lois de distribution florale dans la zone alpine. Bull Soc Vaudoise Sci Nat, 38, 69–130.

61. Edgar, R.C. (2004) MUSCLE: multiple sequence alignment with high accuracy and high throughput. Nucleic Acids Res, 32, 1792–1797.

62. Ho, J., Jain, A. and Abbeel, P. (2020) Denoising Diffusion Probabilistic Models. 10.48550/arXiv.2006.11239.

63. Lewy, D. and Mańdziuk, J. (2023) An overview of mixing augmentation methods and augmentation strategies. Artif Intell Rev, 56, 2111–2169.

64. Shorten, C. and Khoshgoftaar, T.M. (2019) A survey on Image Data Augmentation for Deep Learning. J Big Data, 6, 60.

65. Sharma, N.K., Gupta, S., Kumar, A., Kumar, P., Pradhan, U.K. and Shankar, R. (2021) RBPSpot: Learning on appropriate contextual information for RBP binding sites discovery. iScience, 24, 103381.

66. Zhou, T., Yang, L., Lu, Y., Dror, I., Dantas Machado, A.C., Ghane, T., Di Felice, R. and Rohs, R. (2013) DNAshape: a method for the high-throughput prediction of DNA structural features on a genomic scale. Nucleic Acids Res, 41, W56–62.

67. Brown, T.B., Mann, B., Ryder, N., Subbiah, M., Kaplan, J., Dhariwal, P., Neelakantan, A., Shyam, P., Sastry, G., Askell, A., et al. (2020) Language Models are Few-Shot Learners. 10.48550/arXiv.2005.14165.

68. Papineni, K., Roukos, S., Ward, T. and Zhu, W.-J. (2002) BLEU: a method for automatic evaluation of machine translation. In Proceedings of the 40th Annual Meeting on Association for Computational Linguistics, ACL ‘02. Association for Computational Linguistics, USA, pp. 311–318.

69. Goodstein, D.M., Shu, S., Howson, R., Neupane, R., Hayes, R.D., Fazo, J., Mitros, T., Dirks, W., Hellsten, U., Putnam, N., et al. (2012) Phytozome: a comparative platform for green plant genomics. Nucleic Acids Res, 40, D1178–D1186.

70. Gurevich, A., Saveliev, V., Vyahhi, N. and Tesler, G. (2013) QUAST: quality assessment tool for genome assemblies. Bioinformatics, 29, 1072–1075.

71. Jyoti, Ritu, Gupta, S. and Shankar, R. (2024) Comprehensive analysis of computational approaches in plant transcription factors binding regions discovery. Heliyon, 10, e39140.

72. Li, H. (2018) Minimap2: pairwise alignment for nucleotide sequences. Bioinformatics, 34, 3094–3100.

73. Morisse, P., Lecroq, T. and Lefebvre, A. (2021) Long-read error correction: a survey and qualitative comparison. 10.1101/2020.03.06.977975.

74. Marçais, G., DeBlasio, D., Pandey, P. and Kingsford, C. (2019) Locality-sensitive hashing for the edit distance. Bioinformatics, 35, i127–i135.

75. Delahaye, C. and Nicolas, J. (2021) Sequencing DNA with nanopores: Troubles and biases. PLoS One, 16, e0257521.

76. Berlin, K., Koren, S., Chin, C.-S., Drake, J.P., Landolin, J.M. and Phillippy, A.M. (2015) Assembling large genomes with single-molecule sequencing and locality-sensitive hashing. Nat Biotechnol, 33, 623–630.

77. Tyson, J.R., O’Neil, N.J., Jain, M., Olsen, H.E., Hieter, P. and Snutch, T.P. (2018) MinION-based long-read sequencing and assembly extends the Caenorhabditis elegans reference genome. Genome Res., 28, 266–274.

78. Ondov, B.D., Treangen, T.J., Melsted, P., Mallonee, A.B., Bergman, N.H., Koren, S. and Phillippy, A.M. (2016) Mash: fast genome and metagenome distance estimation using MinHash. Genome Biol, 17, 132.

79. Gupta, S. and Shankar, R. (2023) miWords: transformer-based composite deep learning for highly accurate discovery of pre-miRNA regions across plant genomes. Brief Bioinform, 24, bbad088.

80. Gupta, S., Kesarwani, V., Bhati, U., Jyoti and Shankar, R. (2024) PTFSpot: deep co-learning on transcription factors and their binding regions attains impeccable universality in plants. Brief Bioinform, 25, bbae324.

81. Zhai, J., Gokaslan, A., Schiff, Y., Berthel, A., Liu, Z.-Y., Lai, W.-Y., Miller, Z.R., Scheben, A., Stitzer, M.C., Romay, M.C., et al. (2025) Cross-species modeling of plant genomes at single-nucleotide resolution using a pretrained DNA language model. Proceedings of the National Academy of Sciences, 122, e2421738122.

82. Benegas, G., Batra, S.S. and Song, Y.S. (2023) DNA language models are powerful predictors of genome-wide variant effects. Proceedings of the National Academy of Sciences, 120, e2311219120.

83. Zhou, Z., Ji, Y., Li, W., Dutta, P., Davuluri, R. and Liu, H. (2024) DNABERT-2: Efficient Foundation Model and Benchmark For Multi-Species Genome. 10.48550/arXiv.2306.15006.

84. Mendoza-Revilla, J., Trop, E., Gonzalez, L., Roller, M., Dalla-Torre, H., de Almeida, B.P., Richard, G., Caton, J., Lopez Carranza, N., Skwark, M., et al. (2024) A foundational large language model for edible plant genomes. Commun Biol, 7, 835.

85. Fishman, V., Kuratov, Y., Shmelev, A., Petrov, M., Penzar, D., Shepelin, D., Chekanov, N., Kardymon, O. and Burtsev, M. (2025) GENA-LM: a family of open-source foundational DNA language models for long sequences. Nucleic Acids Res, 53, gkae1310.

86. Quang, D. and Xie, X. (2016) DanQ: a hybrid convolutional and recurrent deep neural network for quantifying the function of DNA sequences. Nucleic Acids Res, 44, e107.

87. Morrell, P.L. and Pakhomov, S.V. (2025) Decoding nature’s grammar with DNA language models. Proceedings of the National Academy of Sciences, 122, e2512889122.

88. Oubounyt, M., Louadi, Z., Tayara, H. and Chong, K.T. (2019) DeePromoter: Robust Promoter Predictor Using Deep Learning. Front. Genet., 10.

89. Bhati, U., Sharma, A., Gupta, S., Kumar, A., Pradhan, U.K. and Shankar, R. (2025) Decoding stress specific transcriptional regulation by causality aware Graph-Transformer deep learning. Current Plant Biology, 43, 100521.

90. Chen, Y., Nie, F., Xie, S.-Q., Zheng, Y.-F., Dai, Q., Bray, T., Wang, Y.-X., Xing, J.-F., Huang, Z.-J., Wang, D.-P., et al. (2021) Efficient assembly of nanopore reads via highly accurate and intact error correction. Nat Commun, 12, 60.

91. Liu, Y., Li, Y., Chen, E., Xu, J., Zhang, W., Zeng, X. and Luo, X. (2024) Repeat and haplotype aware error correction in nanopore sequencing reads with DeChat. Commun Biol, 7, 1678.

92. Hu, J., Wang, Z., Sun, Z., Hu, B., Ayoola, A.O., Liang, F., Li, J., Sandoval, J.R., Cooper, D.N., Ye, K., et al. (2024) NextDenovo: an efficient error correction and accurate assembly tool for noisy long reads. Genome Biol, 25, 107.

