## Supplementary Information for "GANGE: Achieving Sequencing Without Sequencing With Diffusion Guided Generative Genomic Transformer"

**Table S1:** List of some published tools which are used at various stages of the sequencing and assembly pipelines.

| Reference | Name | Broad Classification | Developed for Sequencing Technology | Algorithm |
|---|---|---|---|---|
| **1** | **Bonito** | Basecalling | Oxford Nanopore Technology (ONT) Sequencing Reads | multiple TCSConv-BN-ReLU modules: time-channel separable convolutions (TCSConv), a batch normalization (BN) layer, and a ReLU activation function |
| **2** | **Nanopolish** | Genome Polishing | PacBio and ONT Sequencing Reads | Hidden Markov Model (HMM) |
| **3** | **DeepNano** | Basecalling | ONT Sequencing Reads | Deep Recurrent Neural Networks (RNN) |
| **4** | **Chiron** | Basecalling | ONT Sequencing Reads | Convolutional and bidirectional long short-term memory (Bi-LSTM) layers |
| **5** | **HECIL** | Hybrid Error Correction | PacBio and ONT Sequencing Reads | Iterative Learning |
| **6** | **DeepVariant** | Variant Calling | PacBio and ONT Sequencing Reads | Deep Convolutional Neural Network (CNN) |
| **7** | **Hercules** | Hybrid Error Correction | PacBio and ONT Sequencing Reads | HMM |
| **8** | **Nextpolish** | Genome Polishing | PacBio and ONT Sequencing Reads | K-mer score chain (KSC) algorithm |
| **9** | **DeepNano-blitz** | Basecalling | ONT Sequencing Reads | Bidirectional RNN |
| **10** | **nanoreviser** | Basecalling | ONT Sequencing Reads | CNNs and Bi-LSTM |
| **11** | **NECAT** | Error Correction | ONT Sequencing Reads | a two-step progressive method for nanopore read correction. In the first step, NECAT corrects low-error-rate subsequences (LERS), while in the second step, it corrects high-error-rate subsequences (HERS), of the read |
| **12** | **Consent** | Error Correction and Genome Polishing | PacBio and ONT Sequencing Reads | multiple sequence alignment and local de Bruijn graphs |
| **13** | **MultiNanopolish** | Genome Polishing | ONT Sequencing Reads | HMM |

| | | | | |
|---|---|---|---|---|
| **14** | **Fast-Bonito** | Basecalling | ONT Sequencing Reads | One Convolutional Layer followed by three stacked bidirectional gated recurrent unit (Bi-GRU) layers |
| **15** | **GapPredict** | Gap Filling | Genome Gap Filling | LSTM language model |
| **16** | **deepconsensus** | Error Correction | PacBio CCS Sequencing Reads | Encoder-only Transformer (EoT) |
| **17** | **RNNHC** | Hybrid Error Correction | PacBio and ONT Sequencing Reads | RNN |
| **18** | **Nextpolish2** | Genome Polishing | PacBio HiFi Sequencing Reads | KSC algorithm |
| **19** | **Nextdenovo** | Error Correction and Assembly | PacBio and ONT Sequencing Reads | KSC algorithm |
| **20** | **BigDec** | Error Correction | Illumina Sequencing Short Reads | k-mer spectrum method |
| **21** | **nmTHC** | Hybrid Error Correction | PacBio and ONT Sequencing Reads | generative neural machine translation model with transfer learning |
| **22** | **DeepCorr** | Hybrid Error Correction | PacBio and ONT Sequencing Reads | RNN |
| **23** | **DeepPolisher** | Genome Polishing | PacBio HiFi Sequencing Reads | EoT |
| **24** | **Herro** | Error Correction | ONT Sequencing Reads | CNN and EoT |
| **25** | **deChat** | Error Correction | ONT Sequencing Reads | leveraging the strengths of both de Bruijn graphs and variant-aware multiple sequence alignment to create a synergistic approach |
| **26** | **Goldpolish-target** | Genome Polishing | PacBio and ONT Sequencing Reads | long-read adaptation of the ntEdit+Sealer |
| **27** | **HALE** | Error Correction | PacBio and ONT Sequencing Reads | Minimum Error Correction Framework |
| **28** | **DL-Gapgfilling** | Gap Filling | Plant Genome Gap Filling | CNN layer with a BL-Res layer, and an output layer; BL-Res layer employs stacked Bi-LSTM units with residual connections |

**Table S2: Dataset description of collected promoter-gene pairs.**

| **S.No.** | **Species** | **Instances** |
|---|---|---|
| **1** | *Actinidia chinensis* | 28546 |
| **2** | *Arabidopsis lyrata* | 30000 |

| | | |
|---|---|---|
| **3** | *Amborella trichopoda* | 23332 |
| **4** | *Brassica napus* | 31537 |
| **5** | *Citrus clementina* | 25634 |
| **6** | *Cucumis sativus* | 21619 |
| **7** | *Glycine max* | 35505 |
| **8** | *Hordeum vulgare* | 35806 |
| **9** | *Oryza sativa* | 38773 |
| **10** | *Zea mays* | 34264 |
| **11** | *Arabidopsis thaliana* | 32822 |
| **12** | *Vitis vinifera* | 30000 |
| **13** | *Crocus sativus* | 32000 |
| **14** | *Musa acuminata* | 30000 |
| **15** | *C. elegans* | 21517 |
| **16** | *D. melanogaster* | 17405 |

**Table S3: Details of ONT reads collected from SRA.**

| run_accession | study_accession | secondary_study_accession | tax_id | scientific_name | instrument_platform | instrument_model | library_layout | library_source | read_count |
|---|---|---|---|---|---|---|---|---|---|
| ERR13336040 | PRJEB73710 | ERP158450 | 4530 | Oryza sativa | OXFORD_NANOPORE | PromethION | SINGLE | GENOMIC | 2465104 |
| ERR13336041 | PRJEB73710 | ERP158450 | 4530 | Oryza sativa | OXFORD_NANOPORE | PromethION | SINGLE | GENOMIC | 2784271 |
| SRR16230871 | PRJNA764549 | SRP337810 | 39947 | Oryza sativa Japonica Group | OXFORD_NANOPORE | PromethION | SINGLE | GENOMIC | 3274036 |

| | | | | | | | | | |
|---|---|---|---|---|---|---|---|---|---|
| SRR16611063 | PRJNA775962 | SRP343611 | 3847 | Glycine max | OXFORD_NANOPORE | MinION | SINGLE | GENOMIC | 2493021 |
| SRR29027947 | PRJNA1103102 | SRP503877 | 381124 | Zea mays subsp. mays | OXFORD_NANOPORE | PromethION | SINGLE | GENOMIC | 2354455 |
| SRR29027948 | PRJNA1103102 | SRP503877 | 381124 | Zea mays subsp. mays | OXFORD_NANOPORE | PromethION | SINGLE | GENOMIC | 2047155 |
| SRR29563912 | PRJNA967725 | SRP516277 | 112509 | Hordeum vulgare subsp. vulgare | OXFORD_NANOPORE | MinION | SINGLE | GENOMIC | 534293 |
| SRR29563913 | PRJNA967725 | SRP516277 | 112509 | Hordeum vulgare subsp. vulgare | OXFORD_NANOPORE | MinION | SINGLE | GENOMIC | 936293 |
| SRR30315126 | PRJNA732724 | SRP321352 | 3702 | Arabidopsis thaliana | OXFORD_NANOPORE | PromethION | SINGLE | GENOMIC | 373850 |
| SRR34291213 | PRJNA1284003 | SRP595937 | 9606 | Homo sapiens | OXFORD_NANOPORE | PromethION | SINGLE | GENOMIC | 7096025 |
| SRR34853577 | PRJNA1298774 | SRP605703 | 3847 | Glycine max | OXFORD_NANOPORE | GridION | SINGLE | GENOMIC | 292984 |

**Table S4: Repeat content of species.**

| S.No. | Species | Common Name | Genome Size | Repeat Content |
|---|---|---|---|---|
| **1** | *A. chinensis* | Kiwifruit | ~653 Mb | ~43% |
| **2** | *A. lyrata* | Rock cress | ~207 Mb | ~34% |
| **3** | *A. trichopoda* | Amborella | ~870 Mb | ~48–50% |
| **4** | *B. napus* | Rapeseed | ~1,130 Mb | ~72–75% |

| S.No. | Species | Common Name | Genome Size | Repeat Content |
|---|---|---|---|---|
| **5** | *C. clementina* | Clementine | ~301 Mb | ~20% |
| **6** | *C. sativus* | Cucumber | ~367 Mb | ~24% |
| **7** | *G. max* | Soybean | ~978 Mb | ~59% |
| **8** | *H. vulgare* | Barley | ~5,100 Mb | >80% |
| **9** | *O. sativa* | Rice | ~389 Mb | ~35% |
| **10** | *Z. mays* | Maize | ~2,300 Mb | ~85% |
| **11** | *A. thaliana* | Thale cress | ~135 Mb | ~14% |
| **12** | *V. vinifera* | Grapevine | ~487 Mb | ~41% |
| **13** | *Crocus sativus* | Saffron | ~3.45–4.77 Gb | ~65.57% |
| **14** | *Musa acuminata* | Banana | ~523 Mb | ~30% |

**Table S5: Evaluation of MinHash clustering performance across Jaccard similarity thresholds using 6-mer shingling on ONT read data.**

| Jaccard Threshold | Mean Cluster Size (reads) | Cluster Purity (%) | Consensus Indel Error Rate (%) |
|---|---|---|---|
| 0.4 | 312 ± 87 | 61.3 | 13.4 |
| 0.5 | 187 ± 54 | 74.8 | 11.6 |
| 0.6 | 98 ± 31 | 84.2 | 8.7 |
| 0.7 | 38 ± 12 | 93.6 | 6.7 |
| 0.8 | 14 ± 6 | 97.1 | 7.3 |

Note: Cluster purity is defined as the proportion of reads within a cluster originating from the same genomic region as verified by reference alignment. Consensus indel error rate was computed at 30× genome coverage. The paradoxical increase in consensus indel error rate at threshold 0.8 reflects insufficient per-cluster read depth falling below the 30× minimum.

**Table S6: Evaluation of terminal overlap length on cluster merging performance and downstream consensus accuracy at a fixed Jaccard similarity threshold of 0.7.**

| Terminal Overlap Length (bases) | Mean Cluster Size After Merging (reads) | Cluster Merge Rate (%) | Spurious Merge Rate (%) | Consensus Indel Error Rate (%) |
|---|---|---|---|---|
| 16 | 89 ± 34 | 94.7 | 31.2 | 14.2 |
| 32 | 61 ± 22 | 87.3 | 18.6 | 11.8 |
| 64 | 38 ± 12 | 81.4 | 4.3 | 6.7 |
| 128 | 21 ± 9 | 68.2 | 2.1 | 5.8 |
| 256 | 11 ± 5 | 47.6 | 0.8 | 6.2 |

Note: All evaluations were performed at a fixed Jaccard similarity threshold of 0.7 and 30× genome coverage. Cluster merge rate represents the percentage of initially separate clusters successfully merged. Spurious merge rate is the proportion of merges in which reads from distinct non-overlapping loci were erroneously consolidated, as verified by reference alignment. The increase in spurious merge rate at shorter overlap lengths is attributable to the higher probability of short terminal sequences matching by chance in repeat-dense regions.

**Table S7: List of hyperparameter considered for the optimization of the DDPM model.**

| **Hyperparameter** | **Tried** | **Selected** |
|---|---|---|
| Loss function | Mean Squared Error, Mean Absolute Error, Huber, Cross-Entropy, Binary Cross-Entropy, Categorical Cross-Entropy, Sparse Categorical Cross-Entropy, Kullback-Leibler Divergence, Hinge | Cross-entropy loss function |
| Optimizer | Adadelta, Adagrad, Adam, Adamax, Ftrl, Nadam, RMSprop, SGD, Adam, AdamW, Swish | Swish |
| Learning rate | 0.1-1e-10 | 0.0002 |
| Channel multiplier | 2,4,6,8,....16 | 1,2,2,4 |
| Batch size | 2,4,6,8,....64 | 12 |
| Attention blocks | 2,4,6,8,....16 | 8 |
| Residual blocks | 2,4,6,8,....16 | 8 |

| | | |
|---|---|---|
| Weight decay | 0.1-1e-10 | 0.0005 |
| Beta start | 0.1-1e-10 | 0.0001 |
| Norm groups | 2,4,6,8,....16 | 8 |
| Beta end | 0.1-1e-10 | 0.02 |
| Clip max | 4,3,2,1 | 1 |
| Clip min | -1,-2,-3,-4 | -1 |
| First conv channels | 2,4,6,8,....16 | 8 |

**Table S8: MAE of ten-fold random independent train:test trails of the DDPM model.**

| **Tenfold MAE analysis for DDPM** | | |
|---|---|---|
| **S. No.** | **MAE Training** | **MAE Testing** |
| 1 | 0.323031055 | 0.307952431 |
| 2 | 0.305825142 | 0.307103313 |
| 3 | 0.288002136 | 0.298737317 |
| 4 | 0.295715396 | 0.299823134 |
| 5 | 0.299293075 | 0.302386643 |
| 6 | 0.299478671 | 0.300747847 |
| 7 | 0.292574731 | 0.295089509 |
| 8 | 0.297510944 | 0.299636853 |
| 9 | 0.302997541 | 0.306255857 |
| 10 | 0.296663836 | 0.295776226 |

**Table S9: Accuracy across different number of encoder and decoder layers of the Transformer model.**

| **S.No.** | **Number of Encoder and decoder layers** | **Accuracy** |
|---|---|---|

| | | |
|---|---|---|
| 1 | 1 | 52.62323406 |
| 2 | 2 | 54.92431466 |
| 3 | 3 | 66.37924084 |
| 4 | 4 | 69.14067472 |
| 5 | 5 | 70.60873241 |
| 6 | 6 | 76.28245972 |
| 7 | 7 | 77.86764576 |
| 8 | 8 | 81.37035639 |
| 9 | 9 | 81.73686442 |
| 10 | 10 | 81.9247405 |

**Table S10: List of hyperparameter considered for the optimization of the Transformer model.**

| S.No. | Hyperparameter | Tried | Selected |
|---|---|---|---|
| 1 | Activation function in FFN 1 | Elu, LeakyReLU, relu, selu, sigmoid, softplus, softsign, tanh | Relu |
| 2 | Activation function in FFN 2 | Elu, LeakyReLU, relu, selu, sigmoid, softplus, softsign, tanh | Selu |
| 3 | Optimizer | Adadelta, Adagrad, Adam, Adamax, Ftrl, Nadam, RMSprop, SGD, Adam, AdamW, Swish | Adam |
| 4 | Learning rate | 0.1 - 1e-10 | 0.001 |
| 5 | Weight decay | 0.1 - 1e-10 | 0.00001 |
| 6 | No. of attention head | 1,2,3,4,5,6,7,8,9,10,....64 | 8 |
| 7 | Batch size | 2,4,6,8,....64 | 12 |
| 8 | No. of encoder and decoder | 1,2,3,4,5,6,7,8,9,10 | 8 |
| 9 | No.of neuron in FFN 1 | 8-128 | 42 |
| 10 | No.of neuron in FFN 2 | 8-128 | 64 |

| | | | |
|---|---|---|---|
| 11 | Dropout rate in dropout layer 1 | 0.01-0.5 | 0.12 |
| 12 | Dropout rate in dropout layer 2 | 0.01-0.5 | 0.12 |
| 13 | Dimension Embedding | 8-1024 | 512 |
| 14 | Loss function | Mean Squared Error, Mean Absolute Error, Huber, Cross-Entropy, Binary Cross-Entropy, Categorical Cross-Entropy, Sparse Categorical Cross-Entropy, Kullback-Leibler Divergence, Hinge | Cross-entropy |

**Table S11: MAE of ten-fold random independent train:test trails of the Transformer model.**

| **Tenfold MAE analysis for Transformer** | | |
|---|---|---|
| **S. No.** | **MAE Training** | **MAE Testing** |
| 1 | 0.305390193 | 0.302204017 |
| 2 | 0.3210871 | 0.308687626 |
| 3 | 0.312274841 | 0.313717104 |
| 4 | 0.318566733 | 0.305176936 |
| 5 | 0.311770514 | 0.314371344 |
| 6 | 0.316007129 | 0.316076416 |
| 7 | 0.32696813 | 0.335889034 |
| 8 | 0.31750479 | 0.320084089 |
| 9 | 0.315753937 | 0.317489653 |
| 10 | 0.317197352 | 0.313516575 |

**Table S12: Statistics of the assembly generated with Standard genome protocol and GANGE backed genome assembly.**

| | ***A. thaliana*** | | ***O. sativa*** | |
|---|---|---|---|---|
| **Metrics** | Standard protocol | GANGE backed genome | Standard genome protocol | GANGE backed assembly |

| | | | | |
|---|---|---|---|---|
| | | assembly | | |
| **Number of contigs** | 16 | 9 | 34 | 20 |
| **Total length** | 114.2 | 121.64 | 384.4 | 389.7 |
| **Longest contig length** | 28.6 | 32.12 | 35.6 | 43.2 |
| **misassemblies** | 37 | 29 | 502.03 | 319.16 |
| **indel rate per 100 kbp** | 472.93 | 270.98 | 57 | 32 |
| **BUSCO** | 91.4 | 96.1 | 91.7 | 96.8 |
| **N50** | 21.36 | 23.2 | 29.8 | 30.9 |
| **Genome fraction (%)** | 97.9 | 98.3 | 97.83 | 98.12 |

**Table S13: Statistics of the assembly generated with Standard genome protocol and GANGE backed genome assembly.**

| | ***Homo sapiens*** | |
|---|---|---|
| **Paramters** | Standard protocol | GANGE backed assembly |
| **Number of contigs** | 2 | 2 |
| **Total length** | 232 | 242 |
| **Longest contig length** | 194 | 229 |
| **indel rate per 100 kbp** | 254.12 | 210.25 |
| **N50** | 194 | 229 |

**Table S14: Statistics of the assembly generated with different approach backed genome assembly.**

| **Paramters** | **GANGE** | **HERRO** | **NECAT** | **DeChat** | **NextDenovo** |
|---|---|---|---|---|---|
| **Number of contigs** | 9 | 12 | 14 | 12 | 14 |
| **Total length** | 121.64 | 120.42 | 118.65 | 120.38 | 119.89 |
| **Longest contig length** | 32.12 | 31.1 | 30.3 | 29.7 | 31.9 |

| | | | | | |
|---|---|---|---|---|---|
| **indel rate per 100 kbp** | 270.98 | 287 | 305.93 | 297.23 | 306.24 |
| **N50** | 23.2 | 22.85 | 22.74 | 22.82 | 22.67 |
| **Genome fraction (%)** | 98.3 | 97.2 | 96.5 | 96.1 | 96.8 |

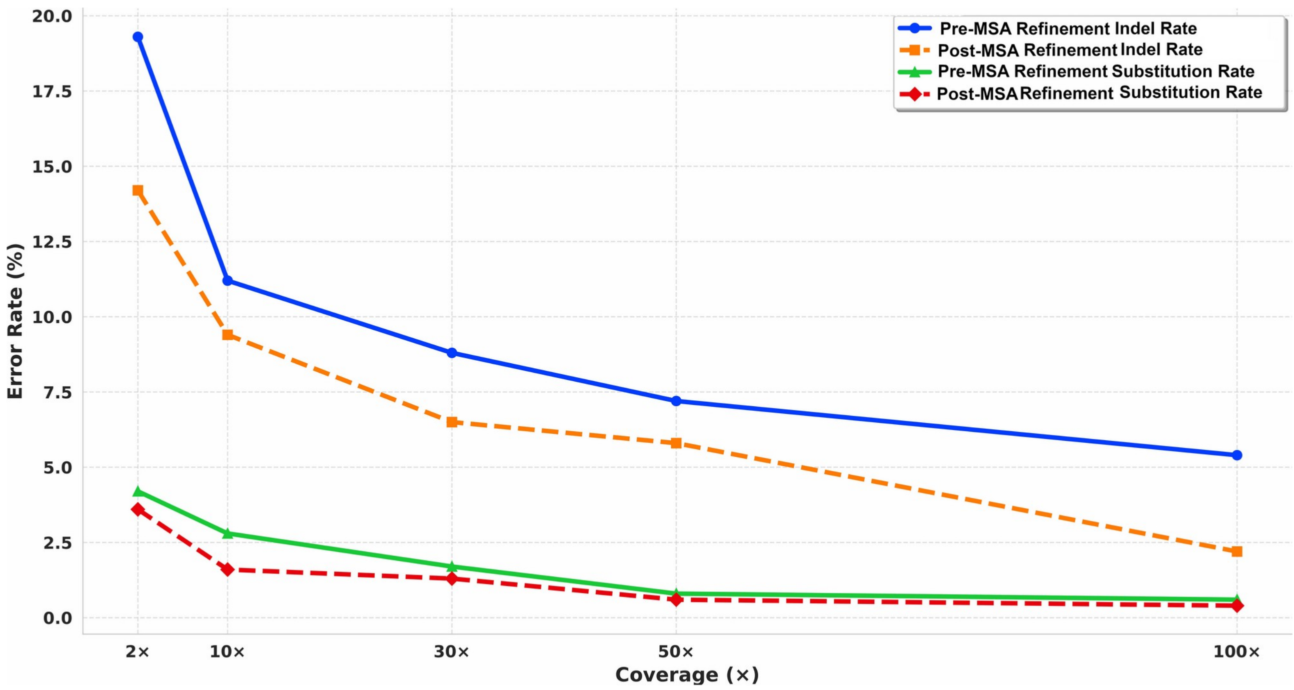


**Figure S1: Error rate on different depths of sequencing coverage before and after MSA refinement.** Plot depicting the relationship between sequencing coverage depth (X) and error rate (%). Error rates decline sharply with increasing coverage from 1X to 50X, followed by a gradual stabilization beyond 50X coverage. This inverse relationship demonstrates that higher sequencing depth improves consensus accuracy by enabling better discrimination between true biological variants and sequencing artifacts.

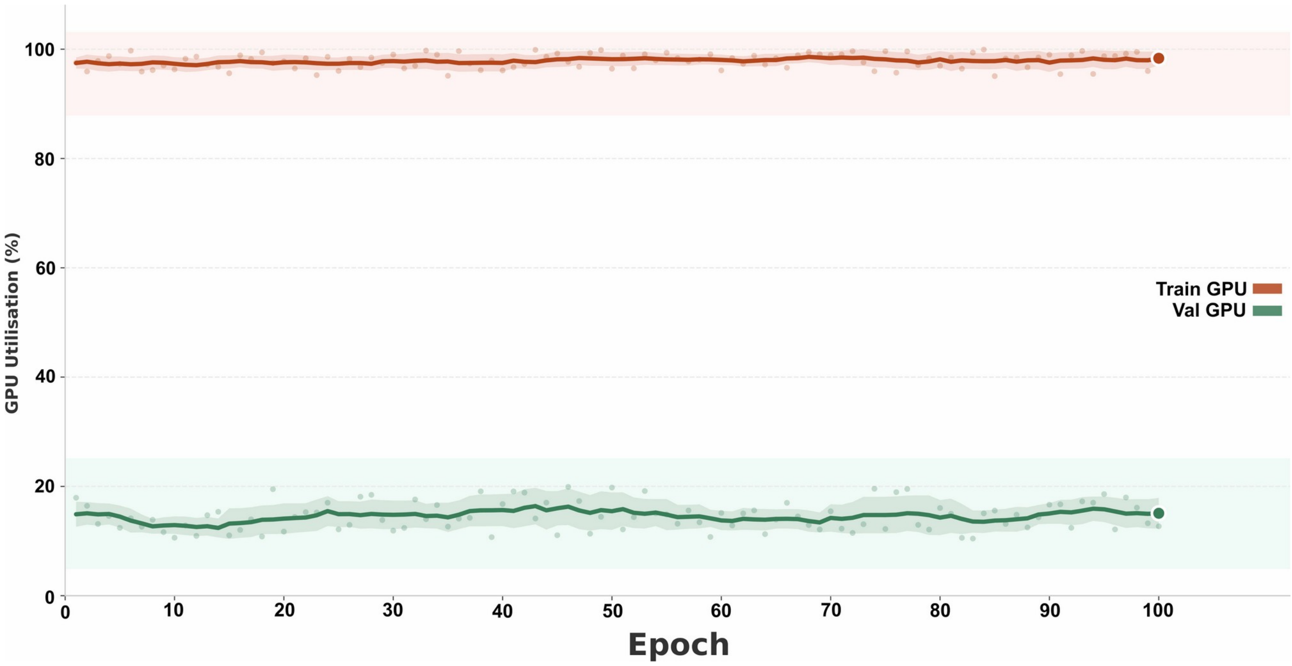


**Figure S2: GPU utilization during training and validation phases of the Transformer system.** Training maintains consistently high GPU utilization (~95-99%), while testing requires substantially lower requirement (~10–20%), confirming frugal application overhead.

**References**

1. nanoporetech/bonito (2026). https://github.com/nanoporetech/bonito
2. Loman,N.J., Quick,J. and Simpson,J.T. (2015) A complete bacterial genome assembled de novo using only nanopore sequencing data. Nat Methods, 12, 733–735.
3. Boža,V., Brejová,B. and Vinař,T. (2017) DeepNano: Deep recurrent neural networks for base calling in MinION nanopore reads. PLoS One, 12, e0178751.
4. Teng,H., Cao,M.D., Hall,M.B., Duarte,T., Wang,S. and Coin,L.J.M. (2018) Chiron: translating nanopore raw signal directly into nucleotide sequence using deep learning. Gigascience, 7, giy037.
5. Choudhury,O., Chakrabarty,A. and Emrich,S.J. (2018) HECIL: A Hybrid Error Correction Algorithm for Long Reads with Iterative Learning. Sci Rep, 8, 9936.
6. Poplin,R., Chang,P.-C., Alexander,D., Schwartz,S., Colthurst,T., Ku,A., Newburger,D., Dijamco,J., Nguyen,N., Afshar,P.T., et al. (2018) A universal SNP and small-indel variant caller using deep neural networks. Nat Biotechnol, 36, 983–987.
7. Firtina,C., Bar-Joseph,Z., Alkan,C. and Cicek,A.E. (2018) Hercules: a profile HMM-based hybrid error correction algorithm for long reads. Nucleic Acids Res, 46, e125.
8. Hu,J., Fan,J., Sun,Z. and Liu,S. (2020) NextPolish: a fast and efficient genome polishing tool for long-read assembly. Bioinformatics, 36, 2253–2255.
9. Boža,V., Perešíni,P., Brejová,B. and Vinař,T. (2020) DeepNano-blitz: a fast base caller for MinION nanopore sequencers. Bioinformatics, 36, 4191–4192.
10. Wang,L., Qu,L., Yang,L., Wang,Y. and Zhu,H. (2020) NanoReviser: An Error-Correction Tool for Nanopore Sequencing Based on a Deep Learning Algorithm. Front. Genet., 11.

11. Chen,Y., Nie,F., Xie,S.-Q., Zheng,Y.-F., Dai,Q., Bray,T., Wang,Y.-X., Xing,J.-F., Huang,Z.-J., Wang,D.-P., et al. (2021) Efficient assembly of nanopore reads via highly accurate and intact error correction. Nat Commun, 12, 60.
12. Morisse,P., Marchet,C., Limasset,A., Lecroq,T. and Lefebvre,A. (2021) Scalable long read self-correction and assembly polishing with multiple sequence alignment. Sci Rep, 11, 761.
13. Hu,K., Huang,N., Zou,Y., Liao,X. and Wang,J. (2021) MultiNanopolish: refined grouping method for reducing redundant calculations in Nanopolish. Bioinformatics, 37, 2757–2760.
14. Xu,Z., Mai,Y., Liu,D., He,W., Lin,X., Xu,C., Zhang,L., Meng,X., Mafofo,J., Zaher,W.A., et al. (2021) Fast-bonito: A faster deep learning based basecaller for nanopore sequencing. Artificial Intelligence in the Life Sciences, 1, 100011.
15. Chen,E., Chu,J., Zhang,J., Warren,R.L. and Birol,I. (2021) GapPredict – A Language Model for Resolving Gaps in Draft Genome Assemblies. IEEE/ACM Trans Comput Biol Bioinform, 18, 2802–2808.
16. Baid,G., Cook,D.E., Shafin,K., Yun,T., Llinares-López,F., Berthet,Q., Belyaeva,A., Töpfer,A., Wenger,A.M., Rowell,W.J., et al. (2023) DeepConsensus improves the accuracy of sequences with a gap-aware sequence transformer. Nat Biotechnol, 41, 232–238.
17. Wang,R. and Chen,J. (2023) RNNHC: A hybrid error correction algorithm for long reads based on Recurrent Neural Network. 10.21203/rs.3.rs-3309460/v1.
18. Hu,J., Wang,Z., Liang,F., Liu,S.-L., Ye,K. and Wang,D.-P. (2024) NextPolish2: A Repeat-aware Polishing Tool for Genomes Assembled Using HiFi Long Reads. genom. proteom. bioinform., 22, qzad009.
19. Hu,J., Wang,Z., Sun,Z., Hu,B., Ayoola,A.O., Liang,F., Li,J., Sandoval,J.R., Cooper,D.N., Ye,K., et al. (2024) NextDenovo: an efficient error correction and accurate assembly tool for noisy long reads. Genome Biol, 25, 107.
20. Expósito,R.R. and González-Domínguez,J. (2024) BigDEC: A multi-algorithm Big Data tool based on the k-mer spectrum method for scalable short-read error correction. Future Generation Computer Systems, 154, 314–329.
21. Wang,R. and Chen,J. (2024) NmTHC: a hybrid error correction method based on a generative neural machine translation model with transfer learning. BMC Genomics, 25, 573.
22. Wang,R. and Chen,J. (2024) DeepCorr: a novel error correction method for 3GS long reads based on deep learning. PeerJ Comput. Sci., 10, e2160.
23. Mastoras,M., Asri,M., Brambrink,L., Hebbar,P., Kolesnikov,A., Cook,D.E., Nattestad,M., Lucas,J., Won,T.S., Chang,P.-C., et al. (2024) Highly accurate assembly polishing with DeepPolisher. bioRxiv, 10.1101/2024.09.17.613505.
24. Stanojević,D., Lin,D., Nurk,S., Sessions,P.F. de and Šikić,M. (2024) Telomere-to-Telomere Phased Genome Assembly Using HERRO-Corrected Simplex Nanopore Reads. 10.1101/2024.05.18.594796.
25. Liu,Y., Li,Y., Chen,E., Xu,J., Zhang,W., Zeng,X. and Luo,X. (2024) Repeat and haplotype aware error correction in nanopore sequencing reads with DeChat. Commun Biol, 7, 1678.
26. Zhang,E., Coombe,L., Wong,J., Warren,R.L. and Birol,I. (2025) GoldPolish-target: targeted long-read genome assembly polishing. BMC Bioinformatics, 26, 78.
27. Barak,P., Gibney,D. and Jain,C. Haplotype-aware long-read error correction.
28. Chen,Y., Wang,Z., Wang,G. and Wang,G. (2026) DL-GapFilling: a novel deep learning framework for improved plant genome gap filling. Brief Bioinform, 27, bbag007.
